# Evolution of preferences in structured populations: genes, guns, and culture

**DOI:** 10.1101/408435

**Authors:** Ingela Alger, Jörgen W. Weibull, Laurent Lehmann

## Abstract

During human evolution, individuals interacted mostly within small groups that were connected by limited migration and sometimes by conflicts. Which preferences, if any, will prevail in such scenarios? Building on population biology models of spatially structured populations, and assuming individuals’ preferences to be their private information, we characterize those preferences that, once established, cannot be displaced by alternative preferences. We represent such uninvadable preferences in terms of fitness and in terms of material payoffs. At the fitness level, individuals can be regarded to act as if driven by a mix of self-interest and a Kantian motive that evaluates own behavior in the light of the consequences for own fitness if others adopted this behavior. This Kantian motive is borne out from (genetic or cultural) kin selection. At the material-payoff level, individuals act as if driven in part by self-interest and a Kantian motive (in terms of material payoffs), but also in part by other-regarding preferences towards other group members. This latter motive is borne out of group resource constraints and the risk of conflict with other groups. We show how group size, the migration rate, the risk of group conflicts, and cultural loyalty shape the relative strengths of these motives.

**JEL codes:** A12, A13, B52, C73, D01, D63, D64, D91.

## 1 Introduction

Preferences are fundamental to economic theory.^1^ If preferences are transmitted across generations and if they affect the expected survival and reproduction–the fitness–of their bearer: which preferences are likely to be favored by evolution and which preferences are likely to disappear? Analysis of the long-term evolution of preference distributions can help understand the proximate drivers and motivation of human behavior in social and economic interactions (Hirshleifer, 1977, Bergstrom, 1996, Binmore 1998, Robson, 2001, Newton, 2018, Alger and Weibull, 2019). Here we build on previous work on strategy evolution in structured populations (Lehmann, Alger, and Weibull, 2015) by studying preference evolution in such populations.

For more than a million years, our ancestors most likely lived in groups of hunter-gatherers (probably ranging from 5 to 150 grown-ups), extending beyond the nuclear family (Grueter, Chapais, and Zinner, 2012, Malone, Fuentes, and White, 2012, van Schaik, 2016, Layton, O’Hara, and Bilsborough, 2012). This population structure, whose defining features are small group size and limited migration between groups (i.e., not all individuals migrate), is thus part of the environment of evolutionary adaptation of the human lineage (e.g., van Schaik, 2016). Analysis of the long-term evolution of preferences should thus take such population structure into account. We here do exactly that, and we ask how such structural features as group size, migration rates between groups, and the risk of conflicts between groups, determine the qualitative nature of the preferences that evolution favors. Combining the economics paradigm of utility-maximizing behavior with methods from population genetics, we obtain predictions about the nature of individuals’ preferences and motivations in the canonical model of evolution in structured populations, the so-called island model of migration originally due to Wright (1931, 1943). The model allows us to examine both genetic and cultural transmission of preferences in such structured populations.

The island model is a textbook evolutionary biology model (see, e.g., Cavalli-Sforza and Bodmer, 1971, Frank 1998, Rousset 2004, Hartl and Clark, 2007), which formally captures in a tractable and stylized way the fact that all natural populations, human or otherwise, are structured into small groups (or bands, or villages) connected to each other by limited migration (or dispersal). Limited migration causes limited genetic and/or cultural mixing in the population, and this results in several individuals from the same group possibly having a recent common ancestor. For example, suppose that a genetically transmitted new trait suddenly appears in one individual. In the next generation, multiple carriers of the new trait may coexist in the same group. Hence, the immediate descendants of the initial mutant are more likely to interact with each other than are individuals sampled at random from the whole population. Such assortative matching, induced by limited migration, even when the mutant trait is rare in the population at large, tends to favor mutant behaviors that promote the survival and/or reproductive success of others in their group. The reason is that such behavior is more likely to benefit other mutants than it would be if all offspring always migrated and matching therefore would be uniformly random (Hamilton, 1964, 1971, Grafen, 1985, Frank, 1998, Rousset 2004). This is the so-called mechanism of *kin selection* in evolutionary biology (Maynard Smith, 1964).^2^ In the biology literature, assortative matching between pairs of individuals is usually quantified by the *coefficient of relatedness*—which indicates the likelihood that interacting individuals share a common ancestor—a quantity that depends on such features of the population structure as group size and migration rates.

By the same token, however, individuals who share a local common ancestor are also more likely to expose each other to fitness externalities, than are randomly selected individuals from the overall population. Indeed, through the local interactions, which occur in islands of finite size, related individuals may harm or enhance each other’s fitnesses (think of young siblings fighting over candy, or individuals teaming up to fight off a common enemy), and such externalities have an impact on selected traits (Hamilton, 1971, Schaffer, 1988, Frank, 1998, Rousset 2004). As assortative matching and local fitness externalities can, in general, not be separated, their joint effects need to be taken into account in order to understand the evolutionary success of traits under limited dispersal, a question that has received much attention in the evolutionary biology literature (see e.g. Hamilton, 1967, and Taylor, 1992,a,b, for pioneering and paradigmatic examples, and Frank, 1998, and Rousset, 2004, for general theoretical treatments).

While clearly relevant for the understanding the evolution of traits relevant in human social interactions, the evolutionary biology literature is yet of limited direct value for economists, because in the bulk of these analyses: (a) the focus is on the evolution of strategies, not preferences, (b) predictions are derived at the level of basic fitness components, such as reproduction and survival, and not at the level of the material payoffs obtained in strategic interactions, (c) transmission is genetic instead of cultural, while cultural evolution is also relevant for the understanding of human behavior. Our model enriches the analysis in all of these dimensions.

We propose a framework in line with that of economists and game theorists, and model the following thought experiment that takes place in a large population over an infinite sequence of demographic time periods. The population is structured into a large number of groups or islands of equal size. Within each group or island, individuals engage in a strategic interaction in which all individuals’ strategy choices may affect the material payoffs to all participants. The strategic interaction is modelled as a game in material payoffs, and the game may be arbitrarily complex and may take place over many stages within each demographic time period. By material payoff we mean a one-dimensional summary measure, like income (or calories). The expected material payoffs, realized in a demographic time period, in turn determine the fitness of each individual in the population in that demographic time period. An individual’s fitness is defined as the expected number of individuals in the following time period who have acquired their trait from him or her. If transmission is genetic, an individual’s fitness is the number of his surviving offspring and the individual himself if he survives. If transmission is cultural, an individual’s fitness is the number of individuals in the next time period who acquired their cultural trait from this individual. Offspring may migrate to other groups or islands, or stay in their natal group or island. Many different transmission scenarios are covered by this model framework. For instance, generations may or may not be overlapping, islands may wage wars against each other, traits may be transmitted culturally from parent to child or by imitation of materially successful individuals, etc.

In all our scenarios, genetic and cultural, the population is initially homogenous; all individuals are *ex ante* identical. Suddenly, a different, mutant heritable trait spontaneously appears in exactly one individual. The original, resident trait is uninvadable if there exists no mutant trait, such that the initial mutant produces enough descendants for its trait to be maintained in the population in the long run.

To study preference evolution, we let the heritable traits be continuous utility functions, defined over all strategy profiles that are possible in the material game that represent the interaction on each island. Together with the individual’s (probabilistic) belief about other group members’ strategy choices, an individual’s utility function guides his or her choice of strategy in the local interaction. We evaluate a utility function’s fitness consequences for its carriers in terms of the expected material payoffs that result in all (Bayesian) Nash equilibria under incomplete information, that is, when each individual’s utility function is his or her private information, but individuals’ beliefs about each others’ strategies are consistent with some (Bayesian) Nash equilibrium. We ask if there exist utility functions that are uninvadable in the sense that any mutant utility function does worse, in terms of its carriers’ expected material payoffs, than the residents, in *all* equilibria. Thus bridging the gap between economics and biology, we obtain links between preferences, material incentives, and population structure (including migration and potential group conflicts). The following four main results emerge from our analysis.

First, we obtain a necessary and sufficient condition for a utility function to be uninvadable. This characterization says that a utility function is uninvadable if and only if all strategies used in any Nash equilibrium among individuals with this utility function are, when viewed as heritable strategies, uninvadable by other strategies.

Second, we identify a class of utility functions that, for any given game in material payoffs, contains an uninvadable utility function. Each utility function in this class can be interpreted as a mix of self-interest and a Kantian concern, both expressed at the fitness level. Specifically, the Kantian concern, driven by kin selection, consists in evaluating one’s behavior in the light of what one’s own fitness would be if others in one’s group were to behave in the same way. This concern vanishes under unlimited migration (that is, when all offspring always migrate) and when groups are very large.

Third, when material payoffs only have marginal effects on fitnesses (a property which arguably holds for many human interactions), uninvadable preferences generically involve a mix of self-interest, a Kantian concern, and also a concern for neighbors, all concerns being expressed at the level of material payoffs. The weight given to the Kantian motive is then proportional to the coefficient of relatedness, but it also depends on fitness externalities between neighbors. The weight on other group members’ material payoffs may be negative (“spite”) or positive (“altruism”), and it depends on the *coefficient of fitness interdependence*, which measures the effect on own fitness that an individual obtains relative to his neighbors by diminishing or enhancing their material payoffs.

Finally, we provide sufficient conditions for the uninvadability of preferences of a particularly simple form, namely, a convex combination of own material payoff and the own material payoff that would arise should all others choose the same behavior. Under these specific conditions, the weight given to the second, Kantian, component is determined by the *coefficient of scaled relatedness*, a coefficient that combines the (standard) coefficient of relatedness with the coefficient of fitness interdependence. This weight allows to determine whether, on balance, equilibrium behaviors are pro-or anti-social, in the sense that equilibrium material payoffs are higher or lower than under selfishness. We show that an increased risk for group conflicts make preferences less anti-social, and, at a critical level of the risk of group conflict, preferences are neither anti- nor pro-social, while at higher risk levels, preferences turn pro-social. Hence, at this intermediate risk of conflict, preferences have only a self-interested and a Kantian component, while at lower (higher) risks, a third component appears, a component that expresses envy or spite if the risk is low, and empathy or altruism if the risk is high. We also show that cultural transmission of preferences may trigger anti-sociality because of local competition for proselytes.

Compared to the existing economics literature on preference evolution in social interactions (see Alger and Weibull, 2019, for a recent survey), our model makes two key innovations.^3^ First, it explicitly analyzes the effects of population structure and limited dispersal upon behavior and preferences. While Alger and Weibull (2013, 2016) investigated the evolutionary stability of preferences under incomplete information, they did so in an abstract model of assortative matching which did not explicitly account for the demographics and population dynamics.^4^ They found that preferences expressing a certain combination of self-interest and a Kantian concern are evolutionarily stable, and that preferences that are behaviorally distinct from these are evolutionarily unstable. They also showed how the weight given to the Kantian concern depends on the assortativity in group formation. While assortativity in those models is treated as an abstract primitive, it here arises explicitly and endogenously from the population structure; group size, rates of survival, migration, and conflicts together determine the probability that rare mutants get to interact with each other–i.e., relatedness. The present model thus contributes to this strand of literature by explicitly modeling the population structure and how it gives rise to assortativity.

Second, it establishes a clear distinction between preferences at the fitness level and preferences at the material payoff level. In the existing economics literature on preference evolution, these are taken to coincide. The model makes it clear that, when preferences are expressed at the level of material payoffs, relatedness must go hand in hand with local fitness interdependence, a force which does not appear in Alger and Weibull (2013, 2016). We here also show how relatedness and fitness interdependence can be formally traced back to group size and limited migration. While we already made this distinction in Lehmann, Alger, and Weibull (2015), we then did not analyze preference evolution. Instead, we asked under what conditions, if any, evolving strategies can be interpreted as chosen by rational individuals endowed with specific utility functions (we examined three candidate utility functions, two of which are described above). The value added of the present paper is that we here analyze preference evolution, rather than strategy evolution, in group-structured populations. In addition, we (a) examine other utility functions than those used to establish the “as if” results in Lehmann, Alger, and Weibull (2015), (b) obtain new results concerning fitness interdependence and scaled relatedness, and (c) analyze a wider class of evolutionary scenarios.

Apart from our previous work, the most closely related work is by Akçay and van Cleve (2012). They investigated the evolutionary stability of preferences parameterized by scalar traits (in the vein of Heifetz, Shannon and Spiegel, 2007a-b). In addition to focusing on complete rather than incomplete information, their model differs from ours in two broad respects. First, since they focus only on the effects of traits on reproduction under genetic transmission, they do not obtain results for preferences over strategy profiles or material payoffs. Second, they focus only on necessary first-order conditions. These conditions express how many offspring an individual is willing to forgo, at the margin, in order to marginally increase the number of offspring of other group members.

The paper is organized as follows. Section 2 describes the model and provides a characterization of an uninvadable trait. Section 3 presents the analysis. In Section 4 we illustrate our results in three canonical evolutionary scenarios, including genetic and cultural evolution, as well as potential “wars” between groups. Section 5 concludes. Mathematical proofs are provided in an appendix.

## 2 Model

This section presents the building blocks of the analysis—the population structure and individuals’ life-cycles—and defines the evolutionary stability criterion that will be used. It also describes what is novel compared to the existing literature.

### 2.1 Population structure

Consider a countably infinite population, divided into infinitely many identical *islands* (groups, locations, or villages), each of constant size *n*. Evolution takes place perpetually and stochastically over time, and time is divided into *demographic time periods*. Individuals are called children or offspring in their first demographic period, the period in which they are born, and grown-ups or *adults* in all other periods of their life. No age distinction is made between adults. Each demographic time period consists of two phases:

- In *Phase 1*, the *n* adults in each island engage in a social or economic interaction with each other, the same on all islands and at all times. The strategies used determine each individual’s *material payoff*, which we take to be income (expressed, for example, in money or in calories). An individual’s strategy choice in the interaction is assumed to be determined by her *preferences* and her beliefs about the strategies used by her island neighbors. Preferences are inherited in childhood from exactly one adult, the individual’s genetic or cultural parent, and are fixed throughout her life.
- In *Phase 2*, the realized material payoffs determine each adult’s survival, and, in case there are exogenous random shocks to entire groups (e.g., warfare, environmental catastrophies), the adult’s entire group’s survival. Individual and group survival probabilities are assumed to be independent of age. The realized material payoffs also determine each adult’s fecundity, which is its number of *offspring* (where offspring are biological if preferences are coded for genetically, and cultural if they are transmitted by way of a cultural process). Following reproduction, offspring, and only offspring, may migrate from their native island to other islands (and this migration takes place in the period they are born). The migration probability *m* is the same for all offspring at all times, and is strictly positive. Moreover, migration is blind in the sense that any migrant picks a destination in a uniformly random fashion.^5^ After migration and competition among the offspring for securing a place on an island, there are exactly *n* adults in each island (group). Offspring who did not secure a place on an island die.

Phase 1 and 2 taken together determine an adult’s *individual fitness*. This is the expected number of her *immediate descendants*, defined as those adults in the next demographic time period who have inherited their preferences from her. These immediate descendants consist of those of the individual’s (genetic or cultural) offspring who survived, and thus became adults in the next demographic time period, as well as the individual herself if she survived into the next period. We next describe in more detail how the interactions and the ensuing individual fitness are formalized in the subsequent analysis.

### 2.2 The interaction

#### 2.2.1 The material game

The material game is formalized as a symmetric non-cooperative normal-form *n*-player game in which each player has access to the same set of strategies (which may be pure or mixed), *X*, a non-empty compact set in some normed vector space. The expected material payoff^6^ accruing to any (adult) individual *i* ∈ {1, 2, …, *n*} on a given island depends on her own strategy, *x*_*i*_ ∈ *X*, and on the vector ***x***_−*i*_ ∈ *X*^*n*−1^ of strategies used by the others on *i*’s island, her *neighbors*. The material payoff function *π* : *X*^*n*^ → ℝ is continuous, and *π* (*x*_*i*_, ***x***_−*i*_) is invariant under permutation of the components of the vector ***x***_−*i*_ ∈ *X*^*n*−1^.^7^ Such permutation invariance holds if, for example, strategies are real numbers, and an individual’s material payoff depends on her own strategy and either the sum, product, maximum or minimum of her island neighbors’ strategies. The material game may be a simple simultaneous-move game or a multi-stage game in which individuals interact over many stages within the demographic time period.

#### 2.2.2 The subjective game

Every (adult) individual in the population has (personal) preferences over strategy profiles, preferences that can be represented by some continuous utility function.^8^ More precisely, every individual *i* has a complete and transitive preference ordering ⪰_*i*_ over strategy profiles (*x*_*i*_, ***x***_−*i*_) ∈ *X*^*n*^, such that there exists a continuous function *u*_*i*_ : *X*^*n*^ → ℝ satisfying *u*_*i*_ (*x*_*i*_, ***x***_−*i*_) ≥ *u*_*i*_ (*y*_*i*_, ***y***_−*i*_) if and only if (*x*_*i*_, ***x***_−*i*_) ⪰_*i*_ (*y*_*i*_, ***y***_−*i*_). Letting ℱ be the set of continuous functions *f* : *X*^*n*^ → ℝ, each individual has preferences that admit representation by some function *u* ∈ Θ ⊆ ℱ, where Θ is the subset of *preference types*, or simply *types*. Each individual of any given type *u* ∈ Θ chooses her strategy so as to maximize the expected value of her utility function under her probabilistic beliefs about her island neighbors’ strategy choices.

In order to carry out our evolutionary analysis of preferences, we need to evaluate the fitness consequences of preferences, and we will do so when individuals’ strategy choices constitute Nash equilibria in the subjective game. (Which is not to say that we assume or believe that interactions are always in equilibrium. We use Nash equilibrium as a systematic reference point.)

### 2.3 Individual fitness

An individual’s fitness may depend on (a) own material payoff, (b) the material payoffs to the individual’s island neighbors, and (c) the material payoffs in the population at large. Dependence on own material payoff is self-evident. Dependence on neighbors’ material payoffs arises as soon as neighbors’ survival and number of offspring influences the competition that one’s own offspring meet when competing for succession of deceased adults on the native island. Own and others’ material payoffs may also affect one’s island’s success probability in wars with other islands. Dependence on material payoffs in other islands has two sources; migration and potential wars between islands. First, an individual’s offspring face competition with offspring from other islands, both when competing for succession of deceased adults on her native island and on other islands. Second one’s island’s success probability in wars may depend on those islands’ material payoffs.

The individual (or direct) fitness function *w* : ℝ^*n*+1^→ ℝ is assumed to be continuously differentiable. We write *w* (*π*_*i*_, ***π***_−*i*_, *π*^*^) for *i*’s fitness, where *π*_*i*_ ∈ ℝ is own material payoff, ***π***_−*i*_ ∈ ℝ^*n*−1^, is the vector of her neighbors’ material payoffs, and *π*^*^ ∈ ℝ is the average material payoff in the population at large.^9^ We also assume that *w* (*π*_*i*_, ***π***_−*i*_, *π*^*^) is invariant under permutation of the components of the vector ***π***_−*i*_. Owing to the assumption of constant group size, average fitness in the population is always equal to 1. The subsequent analysis further presumes that an individual’s fitness is strictly increasing in her own material payoff, strictly decreasing in the average material payoff in the population at large, and that it may be decreasing or increasing, or non-monotonic in other group members’ material payoffs, but never increase more from a neighbor’s increase in material payoff than from the same increase in own material payoff. Formally:

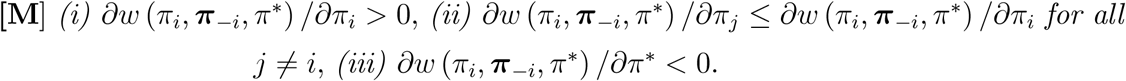

Denote by 𝒫 = ⟨*n, X, π, w*, Θ⟩ a population with (countably) infinitely many islands of size *n*, strategy set *X*, material payoff function *π*, fitness function *w*, and type set Θ ⊆ ℱ. In each demographic time period *t* there is some type distribution *μ*_*t*_ ∈ Δ (Θ) in the population at large. The focus of the analysis is on the dynamics of this type distribution. Analysis will be restricted to type distributions with at most two types present in the population at any given point in time.

Our model adds two novelties to the existing island model literature. First, an individual’s strategy choice is guided by her preferences and beliefs about her island neighbors’ strategy choices. Second, we distinguish material payoffs–here interpreted as income–from individual fitness (see concrete examples of fitness functions in section 5). This distinction is a novelty for the literature on preference evolution in economics, in which fitness is equated with material payoff, and for the evolutionary biology literature, in which income does not appear.

### 2.4 Uninvadability in structured populations

Consider a population 𝒫 which prior to some date *t* = 0 is *homogenous* with some *resident* type *u*. Can this population be invaded by some *mutant* type *v* ≠ *u* that appears at time *t* = 0 in a single adult individual? By “invasion” is meant long-run survival of the mutant type, or, more precisely, that it does not go extinct within finite time. In our model all adults have a positive probability of dying in each period of their lives, and offspring migrate with positive probability, so for a mutant type *v* ≠ *u* to be able to invade the population, it must spread beyond the island where the first mutant appeared. The initial mutant’s descendants may, by way of migration, colonize new islands that were before inhabited exclusively by the resident type *u*. Such colonization, as well as survival and fecundity, depends on all individuals’ material payoffs, which in turn depend on the strategy profiles used in the population. The analysis of a residential type’s invadability or uninvadability is thus an analysis of (non-linear) stochastic population processes involving both (global) demography and (local) strategic interactions.

In order to be able to apply results in the biology literature for stochastic evolution in structured populations to preference evolution, we impose the following *homogeneity assumption* concerning individual’s equilibrium behavior in the subjective game:

> [**H**] On all islands with the same number of mutants, and irrespective of calendar time, the same Nash equilibrium is played, and all residents use the same strategy (say, *x* ∈ *X*), and all mutants use the same strategy (say, *y* ∈ *X*).

As a consequence, on any given island and in any given demographic time period: all residents obtain the same fitness (which in general depends on their strategy, their island neighbors’ strategies, and on strategy profiles in the population at large), and all mutants obtain the same fitness (which likewise depends on their strategy, their island neighbors’ strategies, and on strategy profiles in the population at large). Under these conditions, it is possible to obtain results for the long-run survival, or extinction, of mutants. We proceed in steps towards a definition and characterization of uninvadability.

First, in a population where all individuals are of the same type *u*, all individuals use the same strategy, to be called the *resident strategy*, and this strategy 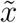 has to satisfy

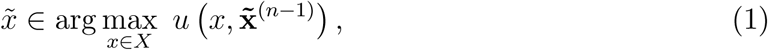

where 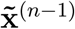 is the (*n* − 1)-dimensional vector whose components all equal 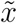. We write *X*_*u*_ for the set of strategies 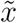 that satisfy (1).^10^ Condition (1) follows from the homogeneity assumption [**H**] and the Nash equilibrium requirement that every individual chooses a strategy that is optimal, given her preferences.

Second, consider a population 𝒫 initially populated by some resident type *u* and in which some strategy in *X*_*u*_ is played by everyone. Let some mutant type *v* appear in exactly one individual at time *t* = 0. Under assumption [**H**], and for any selection of Nash equilibria, one equilibrium for all islands with *k* = 0, 1, …, *n* mutants, respectively, this defines a probability distribution over fitness levels in all future demographic time periods. We define the resident type *u* to be *uninvadable by v* if, for every Nash equilibrium selection, the mutant type *v* goes extinct in finite time with probability one.^11^ A type *u* ∈ Θ is *uninvadable* in Θ if it is uninvadable by all mutant types *v* ∈ Θ.

The notion of *lineage fitness* plays a key role in our characterization of uninvadability. An individual’s *lineage* consists of all of the individual’s descendants, that is, her immediate descendants (her offspring and also herself if she survives), the immediate descendants of her immediate descendants, etc. *ad infinitum*. The individual’s *local lineage* is the subset of her lineage members who live, as adults, in the island where she herself became an adult. Our assumption that the migration rate is positive and constant implies that the random time *T* of first extinction of any individual’s *local lineage* is finite with probability one, and that in time periods before *T* local lineage members may produce emigrants settling on other islands.^12^

Any selection of Nash equilibrium (in the case of complete information, one equilibrium for each number of mutants in a group) defines a Markov chain that induces a unique invariant probability distribution over possible mutant local lineage size realizations (including the realization of the random extinction time *T*), and this occurs irrespectively of whether an island of residents is colonized by a single or several successful mutant emigrants. This probability distribution in turn can be taken to determine the *lineage fitness* of the mutant type *v* given this equilibrium selection, defined as the average *w* fitness of a mutant, the average being taken over (a) all possible local lineage size realizations (each one before the associated random period *T*) and (b) over all possible initial conditions of a local lineage (single or multiple simultaneous emigrant mutants). As long as the mutant is rare in the population, the number of mutants is finite, so the average material payoff earned by individuals of the resident type *u* in those periods is simply *π*^*^ = *π* (*x, x*, …, *x*), where *x* ∈ *X*_*u*_ is the resident strategy in the equilibrium in question.^13^ For any given selection of Nash equilibrium, under assumption [**H**] the lineage fitness of a mutant type *v* ∈ Θ in an otherwise homogeneous population in which all individuals are of type *u* ∈ Θ, can be written in the form

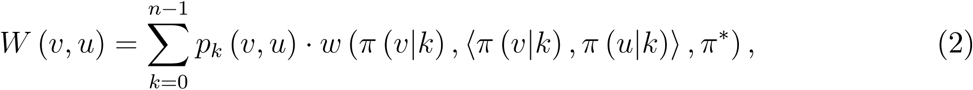

where for each *k* = 0, …, *n* − 1, *p*_*k*_ (*v, u*) is the probability for a mutant uniformly drawn from a local lineage, that *k* = 0, 1, …, *n* − 1 of her neighbors are from this lineage, *π* (*v k*) is the material payoff to the mutant at hand, and ⟨*π* (*v*|*k*), *π* (*u*|*k*)⟩ ∈ ℝ^*n*−1^ is the vector of material payoffs to the mutant’s *n* 1 island neighbors (among whom *k* have the mutant trait *v*, and the other *n* − 1 − *k* have the resident trait *u*). Hence, the lineage fitness of a mutant is the average individual fitness of a representative carrier of the mutant trait. Note that if there are multiple Nash equilibria, there may be several matching probability distributions ***p*** (*v, u*) = (*p*_0_ (*v, u*), …, *p*_*n*−1_ (*v, u*)), one for each selection of Nash equilibrium. Note further that the lineage fitness of the mutant type is well-defined if the mutant type happens to be identical with the resident type; then all individuals in the population have the same lineage fitness, namely *W* (*u, u*) = 1 (since population size is constant).

A positive probability weight *p*_*k*_ (*v, u*) in the definition of *W* for some *k* > 0 means that descendants of the initial mutant face a positive probability of being matched with each other. The overall level of such assortative matching can be usefully quantified by the *coefficient of pairwise relatedness*, defined as

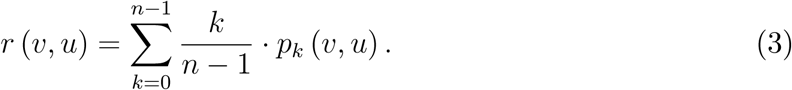

This coefficient measures, for any descendant of the initial mutant of type *v*, the average share of island neighbors who are also descendants of the initial mutant. When migration is complete (*m* = 1, see Section 2.1) or when groups are infinitely large (*n* → ∞), no two group members can be traced back to an initial common ancestor, and thus *p*_*k*_ (*v, u*) = 0 for all *k* > 0, and hence *r* (*v, u*) = 0. But since real-life groups are of finite size, and owing to the cost of dispersal, essentially all natural populations display positive relatedness between group members, i.e., *r* (*v, u*) > 0. This in turn implies that *p*_*k*_ (*v, u*) > 0 for at least some *k* > 0.^14^

We denote by 𝒲 (*v, u*) the set of lineage fitness levels induced by all Nash equilibria compatible with types *v* and *u* in a given population 𝒫. The (potentially empty) set 𝒲 (*v, u*) ⊆ ℝ is compact. Extending the characterization in Lehmann et al. (2016) from types with unique lineage fitness values to types with sets of potential lineage fitness values,^15^ uninvadability can be succinctly characterized as follows: A type *u* ∈ Θ with 𝒲 (*v, u*) ≠ ∅ is *uninvadable* if and only if

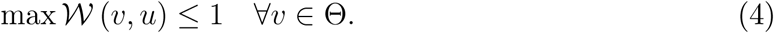

For each mutant type *v* ∈ Θ, this characterization compares the highest possible average lineage fitness of a single initial *v*-mutant, max 𝒲 (*v, u*), with the lineage fitness of any resident individual, *W* (*u, u*) = 1. An uninvadable type *u* thus preempts entry into the population in the sense of obtaining (weakly) higher average lineage fitness that any mutant type can ever obtain.

### 2.5 Nash equilibrium

In order to apply our characterization of uninvadability to preference evolution we need to get a handle on the set of Nash equilibria, which in turn depends on the informational assumptions about the strategic interactions on the islands. We know of three settings that are compatible with homogeneity assumption [**H**], and that admit analysis. In the first setting, each type in the type space Θ ⊂ ℱ has exactly one strategy that it will always use. This is the easiest case, and it can be referred to as “strategy evolution.” In the second setting, all types are permitted, Θ = ℱ, and interactions take place under (maximally) incomplete information, i.e., each individual’s type is his or her private information. In the third setting, interactions take place under complete information, i.e., every individual knows the types of all individuals on her island. Under the homogeneity assumption, each of these settings is amenable to analysis. While one could argue that individuals are likely more knowledgeable about the type distribution in their own island than in the overall population, we nonetheless adopt the incomplete information assumption here, and leave analysis of complete information for future research. The reason is that the incomplete information setting is not only known to provide benchmark results to which results derived under complete information assumptions can be fruitfully compared (for a recent survey, see Alger and Weibull, 2019), but is also likely to be the default case under genetic transmission since information about genotype is generally incomplete (e.g., Frank 1998, chapter 6).

It remains to define the set of Nash equilibria that will be used to calculate any mutant’s lineage fitness under incomplete information. We assume that individuals’ probabilistic beliefs about the type distribution among their neighbors are statistically correct. In particular, every individual of the resident type *u* (correctly) believes that all other individuals on his or her island are (with probability one) of her type, and every mutant (correctly) believes that the types of his or her island neighbors are drawn according to the mutant lineage’s probability distribution ***p*** (*v, u*) = (*p*_0_ (*v, u*), …, *p*_*n*−1_ (*v, u*)). For any given resident strategy 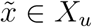, all mutants are, by homogeneity assumption [**H**], assumed to use one and the same strategy, say 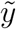 that, moreover, is a best response for them, with their utility function *v*, and given the matching probability distribution that they face:

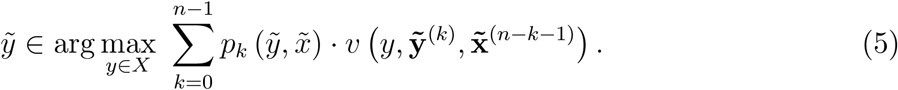

Here 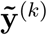 is the strategy vector whose *k* components all are 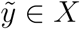, and 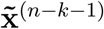 the strategy vector whose *n* − *k* − 1 components all are 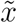, and the matching probabilities are from now on and throughout written directly as a function of the equilibrium strategies played. Given the resident and mutant types, *u, v* ∈ Θ, a strategy pair 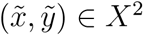 is a (type-homogenous) *Nash equilibrium* if 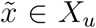 and 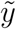 satisfies (5). Let *B*_*NE*_ (*u, v*) ⊆ *X*^2^ denote the set of such Nash equilibria. Any such Nash equilibrium defines all remaining material payoffs *π* (*v*|*k*) and *π* (*u*|*k*) in (2).^16^

## 3 Analysis

It turns out that it is useful, as a first step, to examine preference types which induce commitment to some particular strategy. To be more specific, let Θ ⊂ F consist of all utility functions *u* : *X*^*n*^ → ℝ of the form *u* (*x*_*i*_, ***x***_−*i*_) ≡ ‖*x*_*i*_ − *x*‖^2^ for some *x* ∈ *X*. All individuals with types in this set Θ each have a unique dominant strategy, and we will identify types by their dominant strategy; Θ = *X*. Under such *strategy evolution*, for a resident type *x* and a mutant type *y*, the set 𝒲 (*y, x*) is a singleton, and max 𝒲 (*y, x*) = *W* (*y, x*), where

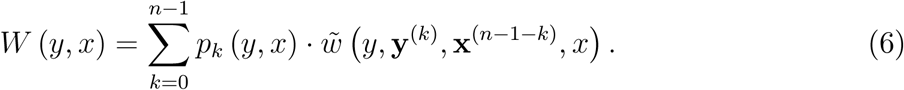

Here 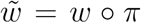 is the composite function which gives the fitness of any individual *i* who plays strategy *x*_*i*_ ∈ *X* when the others on his or her island play ***x***_−*i*_ ∈ *X*^*n*−1^, while some strategy *x*^*^ ∈ *X* is played by all individuals on all other islands:

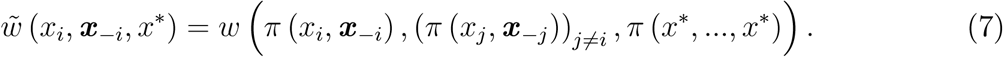

The population size being constant over time, we note that 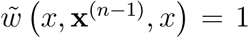 for all *x* ∈ *X*. A necessary and sufficient condition for a strategy *x* ∈ *X* to be uninvadable under strategy evolution is readily obtained by applying condition (4), resulting in:

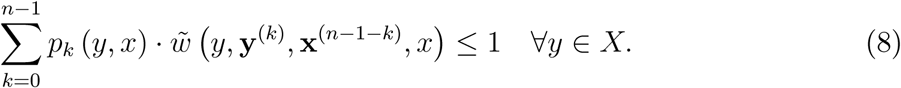

Equivalently, an uninvadable strategy *x* can be seen as preempting entry into the population by earning the maximal lineage fitness that can be obtained in a population where the resident strategy is *x*; that is

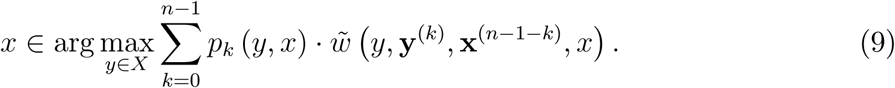

In other words, a strategy is uninvadable if and only if it is a best reply to itself in terms of lineage fitness. We denote by 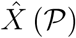 the (potentially empty) set of uninvadable strategies in population 𝒫 = ⟨*n, X, π, w*, Θ⟩.

As a second step we use these observations to write the condition for a type *u* ∈ ℱ to be uninvadable, (4), in an operational form:

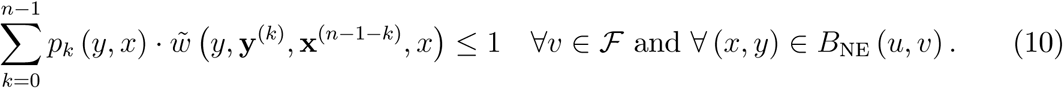

This immediately leads to our first result.

### Proposition 1

*In a population* 𝒫, *a utility function u is uninvadable in* ℱ *if and only if* 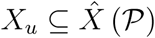.

In other words, for a utility function to be uninvadable, it must induce resident Nash equilibrium strategies that are all uninvadable. However, a utility function does not need to give rise to all uninvadable strategies in resident Nash equilibrium; any strategy in 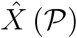 that would not belong to *X*_*u*_ would simply not be played by residents, and would thus not be subject to potential invasion by mutants.^17^

The expression on the left-hand side of (10), however, shows that characterization of uninvadable preferences involves a major challenge, because the matching probabilities may depend both on the resident and mutant strategies, in all time periods when mutants are around.^18^ In the second part of the analysis below, we analyze a model in which independence of the matching probabilities on the strategies played arises endogenously. This model will allow us to fully characterize the set of uninvadable preferences at the level of material payoffs. Prior to that, however, we report results on uninvadable preferences at the level of fitnesses.

### 3.1 Utility and fitness

In spite of the challenge posed by the dependence of the matching probabilities on the strategies played by residents and mutants, we show that one particular class of utility functions stands out, in the sense that there always exists a utility function in this class for which some resident strategy is uninvadable.

#### 3.1.1 The general case

For any given strategy *x*^*^ ∈ *X*, let the utility function *u*_*x\**_ : *X*^*n*^ → ℝ be defined by

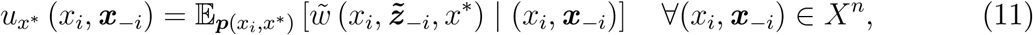

where ***p*** (*x*_*i*_, *x*^*^) = (*p*_0_ (*x*_*i*_, *x*^*^), *p*_1_ (*x*_*i*_, *x*^*^), …, *p*_*n*−1_ (*x*_*i*_, *x*^*^)) is the vector of matching probabilities that would be induced in population 𝒫 if residents played *x*^*^ and mutants played *x*_*i*_. Here 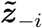 is a *random strategy-profile* such that with probability *p*_*k*_ (*x*_*i*_, *x*^*^) (for each *k* = 0, 1, ..*, n* − 1) exactly *k* of the *n* − 1 components in ***x***_−*i*_ are replaced by *x*_*i*_, with equal probability for each such subset of *k* replaced components, while the remaining components in ***x***_−*i*_ keep their original value. Then:

##### Proposition 2

*Any utility function* 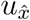 *of the form (11) such that* 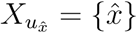 *is uninvadable in ℱ. Moreover, each uninvadable strategy* 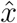 *is also a resident strategy under the utility function* 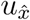.

This proposition identifies a sufficient condition for a utility function of the form (11) to be uninvadable. The condition is that the utility function has a unique resident strategy. Moreover, if the population structure admits multiple uninvadable strategies, then there are multiple utility functions of the form (11) that may be uninvadable, one for each 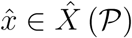. Interestingly, then, different utility functions may arise in different populations with the same population structure. The reason is that the residential strategy affects the matching probabilities (which further explains why this result differs sharply from models in which the assortativity in the matching process is exogenous). There may of course be other uninvadable utility functions than those of the form (11) (see below). Nonetheless, Proposition 2 has a powerful implication: any uninvadable utility function must give rise to a resident strategy that is also a resident strategy under 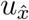 for some 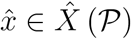.

An individual with the utility function *u*_*x\**_ can be seen as following a probabilistic version of Kant’s categorical imperative (Kant, 1785) at the fitness level; she evaluates the strategies at her disposal in the light of what would happen to her own fitness in the hypothetical scenario in which others would probabilistically use her strategy, according to the probability distribution ***p*** (*x*_*i*_, *x*^*^).^19^ For illustrative purposes, we state *u*_*x\**_ explicitly for *n* = 2 (then calling own strategy *x*_*i*_ and the opponent’s strategy *x*_*j*_) and *n* = 3 (then calling the opponents’ strategies *x*_*j*_ and *x*_*k*_):

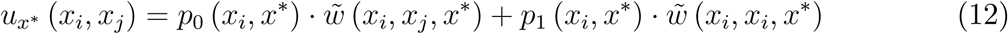

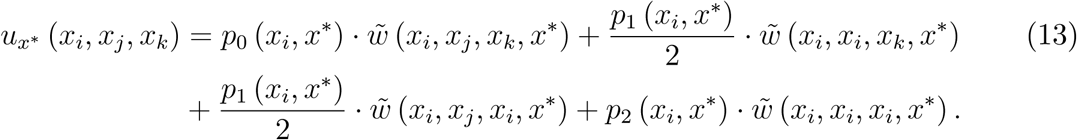

Note that the weights ***p*** (*x*_*i*_, *x*^*^) in the *u*_*x\**_ utility function depend on the individual’s own strategy *x*_*i*_ in the present, whereas in the lineage fitness the matching probabilities depend on the strategy played by mutants individuals living over several (and past) demographic time periods. This highlights the difference between lineage fitness, which is an objective measure, and utility, which is subjective. The dependence of the weights ***p*** (*x*_*i*_, *x*^*^) on own strategy *x*_*i*_, however, questions the operational relevance of *u*_*x\**_ as an analytically and conceptually useful utility function. As such, we now turn to study cases where the matching probabilities in the lineage fitness no longer depend on the mutants’ strategy; this will enable us to turn to utility functions with weights that do not depend on the individual’s strategy.

#### 3.1.2 The differentiable case

Suppose that the following differentiability assumption holds:^20^

[**D**] *(i) X* = ℝ, *(ii) π* : ℝ^*n*^ → ℝ *is continuously differentiable, and (iii) p*_*k*_ : *X*^2^ → [0, 1] is differentiable for all *k* ∈ {0, 1, …, *n* − 1}.

In the next proposition, which states a necessary condition for a strategy 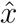 to be uninvadable, 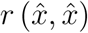 is the coefficient of pairwise relatedness (see (3)) in a population where all individuals play 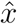, and a subscript *i* on 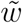 denotes the partial derivative with respect to the *i*-th argument.

##### Proposition 3

*If [****D****] holds and* 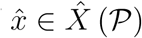, *then*

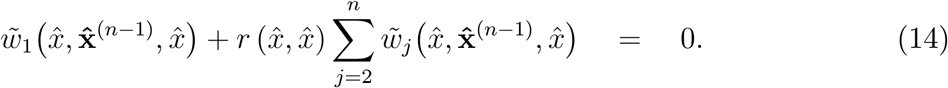

The first term is the marginal fitness benefit of the individual’s own strategy, while the second term is the sum of the marginal fitness benefits conferred by others, weighted by the coefficient of pairwise relatedness. This equation is nothing but the marginal version of Hamilton’s rule (Hamilton, 1964, 1970, Franck, 1998), which provides the necessary first-order condition for an (interior) strategy to be uninvadable (see equation (3) in Taylor and Frank, 1996, or equation (7.5) in Rousset, 2004).^21^ Such first-order conditions are standard in the biology literature, but for the sake of completeness we provide a proof in the appendix.

Consider the utility function defined by

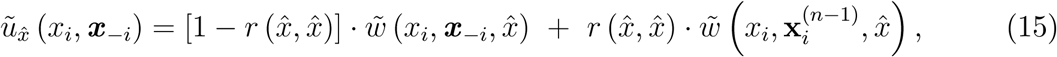

where 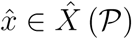, and 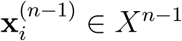 is the strategy vector whose *n* − 1 components all equal *xi*. Clearly, Propositions 1 and 3 together imply that if 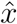 is the unique resident strategy under 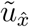, then this utility function is uninvadable.

An individual equipped with the utility function in (15) evaluates her strategy, *x*_*i*_, both in terms of how it affects her own fitness, given the neighbors’ strategies and the strategy played in the population at large, reflected in the first term, and how her strategy *x*_*i*_ would affect her fitness should her neighbors, hypothetically, also use it, reflected in the second term. This is reminiscent of *homo moralis* preferences (see, in particular, Proposition 3 of Alger and Weibull, 2016), although an important difference is that here the utility function in (15) is defined for a certain reference strategy.

### 3.2 Utility and material payoffs

We now turn to an approach in which the matching probabilities still depend on the transmission process but are independent of the strategies used. This approach, in biology called *weak selection* (see, e.g., Nagylaki, 1992, 1993), assumes that fitness effects from the interaction in question are small. Arguably, this approach is highly relevant for the social sciences, since it generates predictions regarding those preferences that guide behaviors in minor everyday interactions, those with only small effects on lifetime fitness.

#### 3.2.1 Weak selection

Formally: for each *x* ∈ *X* and ***y*** ∈ *X*^*n*−1^ let an individual’s material payoff be a convex combination of two terms,

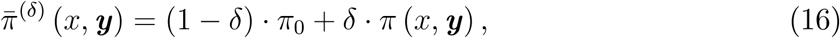

where *π*_0_ is baseline material payoff, assumed identical for all individuals, and *δ* ∈ (0, 1) is the share of the material payoff that emanates from the present material game interaction. This factor *δ* is the *intensity of selection*. Thus, for *δ* ∈ (0, 1) fixed, the fitness of individual *i* is now of the form

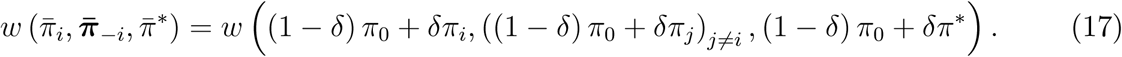

Weak selection amounts to considering the limit as *δ* tends towards 0.^22^ Importantly, under weak selection, the matching probabilities, while still depending on the transmission process, do not depend on the strategies *x* and *y* (for any population 𝒫 = ⟨ *n, X, π, w*, Θ⟩ satisfying assumption [**M**]). The probability for a randomly drawn descendant of an ancestor, be it a resident or mutant, to coexist in its island with *k* other descendants of the same ancestor is then solely determined by the vital rates in a population in which everybody uses the same strategy *x*, no matter which. In biology this is referred to as the *neutral process*. This in turn has profound implications for the ability of a mutant trait to invade, since it means that the strategy played by residents matters only insofar as it affects the local success of mutants.

Let 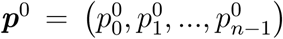 denote the vector of matching probabilities induced by the neutral process. Proposition 2 still holds under weak selection: individuals playing some uninvadable strategy 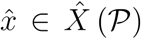 may be viewed as if they were striving to maximize the utility function 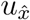, with the matching profile now given by ***p***^0^. This utility function is a sum of individual fitnesses. However, as is shown in the next proposition, under weak selection there is also an uninvadable utility function which describes preferences at the level of material payoffs, and which does not depend on any reference strategy. Let *v*^0^ : *X*^*n*^ → ℝ be defined by

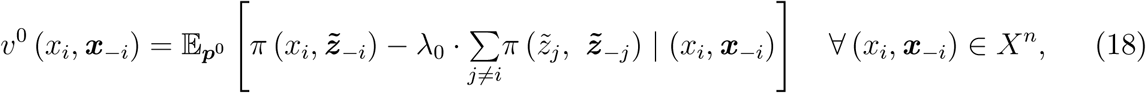

where 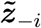 is defined in the same way as in (11), and

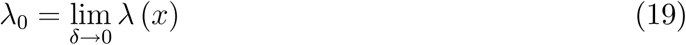

is the *coefficient of fitness interdependence* under weak selection, where

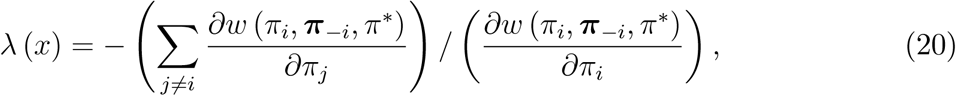

evaluated when all individuals in the population use the same strategy *x* ∈ *X*. Hence, *λ*_0_ measures the marginal effect of neighbors’ material payoffs on own fitness, relative to the marginal effect of own material payoff on own fitness, in a population in which all individuals play the same strategy, and in the limit as *δ* tends to zero. A positive coefficient *λ*_0_ can be interpreted as there being competition for local resources: an increase in the material payoffs to neighbors then reduces an individual’s fitness. A negative coefficient *λ*_0_ means that there is a positive externality at the level of material payoffs between neighbors: an increase in the material payoffs to neighbors then increases an individual’s fitness.

Our next result establishes that selection favors the so-defined utility function, which was used in the context of strategy evolution in Lehmann, Alger, and Weibull (2015), and rejects other utility functions unless they induce an identical best reply to some resident equilibrium for *v*^0^:

##### Proposition 4

*The utility function v*^0^ *is uninvadable in* Θ = ℱ *under weak selection. A utility function u* ∈ Θ *is invadable under weak selection if* 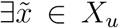 *such that* 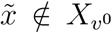. *Moreover*, 1 − *n* ≤ *λ*_0_ ≤ 1.

An individual with the utility function *v*^0^ is but that of the familiar *Homo oeconomicus* if *λ*_0_ = 0 and 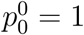. By contrast, if *λ*_0_ ≠ 0 and 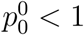, the individual evaluates any strategy profile (*x*_*i*_, ***x***_−*i*_) by pondering his expected *material payoff advantage* over his neighbors, 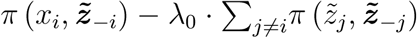, if all, some, or none of the others in her island would use the same strategy as herself (drawn randomly according to *p*^0^), instead of playing their strategies, given by ***x***_−*i*_.

To illustrate how the *v*^0^ goal function is related to preferences studied in behavioral and experimental economics, we briefly consider the two-player case. By writing the utility function as

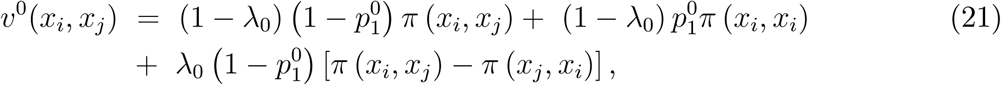

it can be interpreted as the sum of three terms, where the first represents “pure self-interest” (own material payoff), the second a Kantian concern (what is the “right thing to do if others in the population act like me”), and the third a “comparison with the Joneses” (the difference between own material payoff and that of the neighbor). Note also that a positive weight *λ*_0_ > 0 expresses a form of *envy* or *spite*; if instead *λ*_0_ < 0, then it is as if individuals care positively, or *altruistically*, about their neighbors’ material payoffs.

##### Remark 1

*Part of the economics literature on the evolutionary stability of strategies and preferences relies on models in which rare mutants may have a positive probability of being matched with each other, even in the limit as the share of mutants tends to zero (Bergstrom, 2003, Alger and Weibull, 2013, 2016). These limit matching probabilities are taken to be independent of the strategies being played. Hence, they may be interpreted as the vector of matching probabilities* ***p***^0^ *in the neutral process.*

#### 3.2.2 The differentiable case

We finally turn to return to the general model, i.e., selection need not be weak, and consider settings where [**D**] holds. In a population in which all individuals play *x*, let *κ* (*x*) denote the coefficient of scaled relatedness, defined as

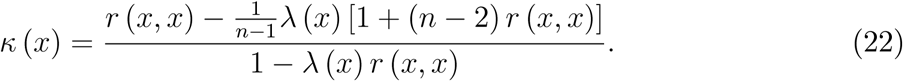

Then we obtain a result that (unlike Proposition 3) is new to the evolutionary biology literature:^23^

##### Proposition 5

*If [****D****] holds and* 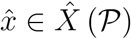, *then*

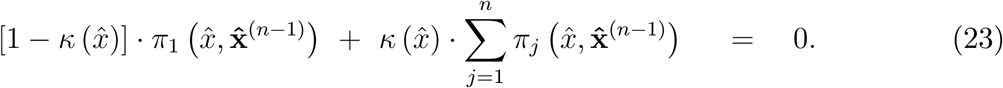

Like 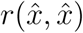, the coefficient 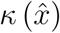 can be interpreted as a marginal substitution rate: it gives the number of units of own material payoff that any given individual is willing to forgo to increase the material payoff of each neighbor by one unit. Absent any fitness interdependence, i.e., if 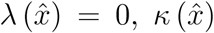 would simply equal relatedness 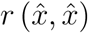. To see exactly how 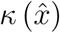 accounts for fitness interdependence, consider first the case when there is but one neighbor, that is *n* = 2. A payoff transfer to this neighbor increases competition from the neighbor at rate 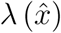 (since 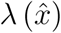 measures the relative increase in competition in the neighborhood of an individual when its payoff is varied, see (20)). The fitness benefit to the donor from giving the transfer to the neighbor is thus reduced by 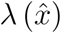, so that the numerator in (22) becomes 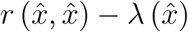. Moreover, a transfer of resources to the neighbor alleviates the competition that the neighbor experiences, and the neighbor is related to the donor according to coefficient 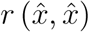. Hence, the cost of the transfer is reduced by 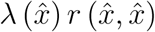, which explains the denominator in (22).

When there are multiple neighbors, *n* > 2, a transfer given to one neighbor enhances the competition by 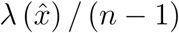, but also for the (*n* − 2) other neighbors, each of which is related to the donor according to coefficient 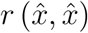. Therefore, the fitness benefit of the transfer to the donor is reduced by 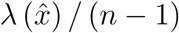 times the term in square brackets in the numerator; which explains the numerator of 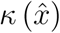. In the denominator, the cost of the transfer is still reduced by 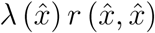, which is the expected alleviation of competition that the transfer induces for the individual’s neighbors (recall that 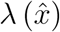 accounts for all neighbors through the term (*n* − 1)).

In view of the necessary first-order condition (23), it may be of interest to consider the utility functions 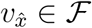 defined by

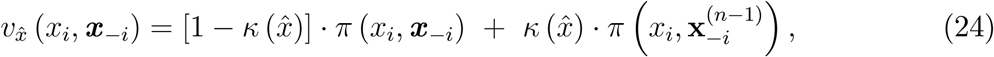

where 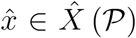. Since (23) implies that 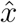 satisfies the necessary first-order condition for an interior symmetric Nash equilibrium of the *n*-player game in which all players have utility function 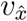, Proposition 1 implies that 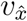 is an uninvadable utility function if 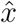 is the unique resident strategy under 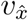.

In sum, in a population in which all individuals play some interior uninvadable strategy 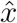, these individuals may (under some conditions) be perceived as having a Kantian concern at the fitness level as well as at the material payoff level. Importantly, the strength of the Kantian (or other-regarding) concern at the fitness level, measured by 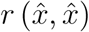, typically differs from the strength of the Kantian (or other-regarding) concern at the material payoff level, measured by 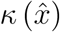, as shown next:

##### Proposition 6

*Suppose that [****D****] holds and that* 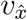 *is uninvadable. The weight* 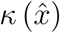 *attached to the neighbors’ material payoffs in the function* 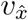 *lies in the interval* [−1, 1]. *Furthermore*, 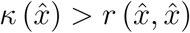 *if and only if* 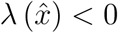.

We note that a necessary and sufficient condition for 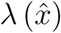 to be negative is that in a population where everybody plays 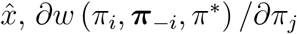 is positive (this partial derivative being the same for all *j* ≠ *i*). We finally note that under weak selection (22) becomes:

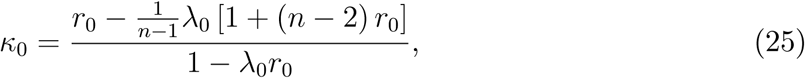

where *λ*_0_ is defined in (19) and *r*_0_ = lim_*δ*→0_ *r* (*x, x*).

## 4 Three canonical scenarios

We have reported general theoretical results on how fitness consequences of material payoffs may be expected to affect preferences over material payoff outcomes. In this section we apply these general results by examining three canonical evolutionary scenarios. For each scenario we calculate the associated coefficients of relatedness *r*, fitness interdependence *λ*, and scaled relatedness *κ* (all the calculations can be found in the appendix). Once these coefficients have been identified, equations (18), (15) and (24) provide closed-form representations of uninvadable utility functions, expressed in terms of the material payoff function that represents the strategic interaction at hand. We note that these coefficients are independent of the material payoff function in question, so the obtained utility representations can be carried over from one material game to any other material game. Also, for all these scenarios the approximate explicit expression for the matching probabilities can be applied (see equation (78) in the Appendix), so the preferences can be fully evaluated in terms of the aforementioned coefficients.

### 4.1 Scenario A: Genes

If types are genetically determined, a possible fitness function is:

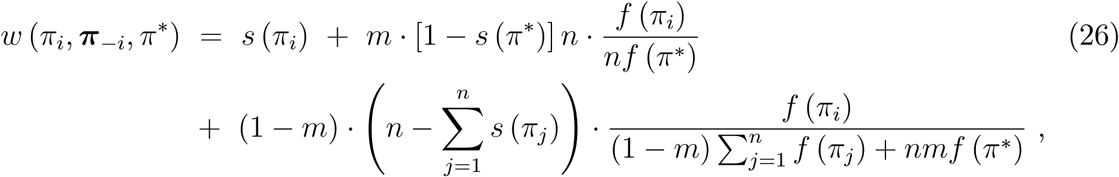

where *s* (*π*_*i*_) ∈ [0, 1] is the probability that *i* survives to the next demographic time period, *f* (*π*_*i*_) > 0 is *i*’s expected number of offspring (who will have inherited *i*’s type), and 0 < *m* ≤ 1 is the probability for each offspring to migrate to another island. These vital events— survival, reproduction and migration–are assumed to be statistically independent. In each island the deceased adults, if any, are replaced by (uniformly) randomly drawn aspiring offspring, native and immigrant. The fortunate ones settle and become adults while the unfortunate ones die. The third term is thus the expected number of *i*’s offspring who manage to secure a “breeding spot” on the natal island. It is the product of three factors: (a) the probability for not migrating, (1 − *m*); (b) the number of available spots on the island; and, for each available spot, (c) the competition for the spot, among native and migrating offspring from other islands, where *f* (*π*^*^) is he fecundity in the population at large. The second term is the expected number of *i*’s offspring who migrate and manage to secure a breeding spot on another island: each offspring who migrates to another island competes against *nf* (*π*^*^) other individuals for the *n* available spots.

#### Remark 2

*For a more detailed derivation of an equation like (26) from the random variables that underlie survival and reproduction, see Lehmann and Balloux (2007). In particular, since the total number of islands is infinite, the probability is zero for the event that more than one of i’s offspring happen to migrate to the same island. Moreover, when the expected number of offspring is large, as we here assume, then the event that there are fewer aspiring offspring than there are available slots in an island is negligible, and the ratios between the expected numbers of offspring, in (26), equal the expectations of the ratios of the underlying random numbers of offspring.*

Considering the case where the survival probability is constant, *s* (*π*_*i*_) = *s*_0_, the coefficient of relatedness equals

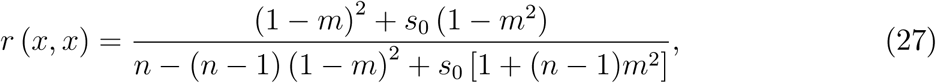

and the coefficient of fitness interdependence equals

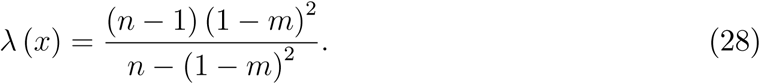

Both coefficients turn out to be independent of the reference strategy *x*. Hence, the utility functions *u*_*x*_ and *v*_*x*_, defined in equations (15) and (24), are independent of what strategy *x* is used in the population at large and can, in this evolutionary scenario, be explicitly parametrized in terms of the migration rate *m*, group size *n*, and survival probability *s*_0_. Both *r* (*x, x*) and *λ* (*x*) are strictly positive for all *n*, all *m* ∈ (0, 1), and all *s*_0_ ∈ (0, 1). By contrast, if *m* = 1, the probability of interacting with an individual from the same lineage is nil, *r* (*x, x*) = 0, and, moreover, there is no fitness benefit from out-competing neighbors materially, *λ* (*x*) = 0. Moreover, both coefficients are decreasing in *m*, and *r* (*x, x*) is increasing in *s*_0_. Substituting (27) and (28) into (22), we obtain:

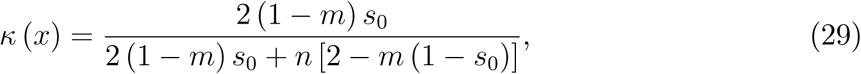

which is strictly positive for any *m* ∈ (0, 1) and *s*_0_ > 0, but nil for *m* = 1 and for *s*_0_ = 0. In other words, in this evolutionary scenario, when *s*_0_ = 0 but *m* ∈ (0, 1), any uninvadable utility function must be as if individuals are pro-social at the level of fitnesses (*r* (*x, x*) > 0), but are purely selfish at the level of material payoffs (*κ* (*x*) = 0). Furthermore, a positive survival probability *s*_0_ > 0 induces pro-sociality (*κ* (*x*) > 0). However, note that *κ* (*x*) is decreasing in island size *n* and in migration rate *m*. In fact, it vanishes as *n* becomes infinitely large. Figure 1 shows how *κ* (*x*) depends on *m* when *s*_0_ = 1*/n*, for *n* = 2 (black solid) and *n* = 10 (black dashed), and when *s*_0_ = 0.8 for *n* = 2 (blue) and *n* = 10 (blue dashed), as well as *s*_0_ = 0 (pink).

**Figure 1:**
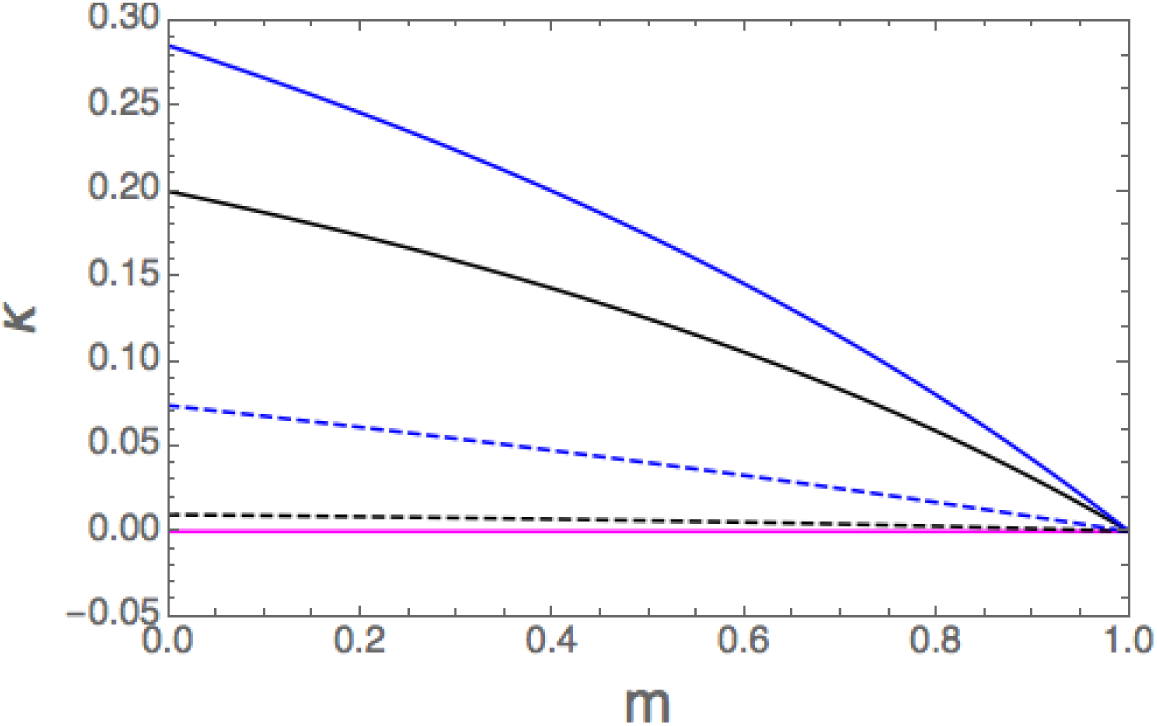
The value of *κ* (*x*) as a function of the migration rate *m*.

#### Remark 3

*In the biology literature, the island model has become a work-horse model to analyze conditions favoring pro- and anti-sociality at the level of survival and reproduction in spatially structured populations. This literature has a well-known result known as Taylor’s cancellation result, a useful yardstick for understanding how changes in the transmission scenario can tip the balance either in the direction of pro-sociality or anti-sociality. Our result that κ* (*x*) = 0 *for s*_0_ = 0 *is in line with this result. When s*_0_ = 0, *(14) boils down to* 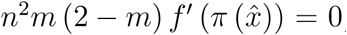, *which implies that* 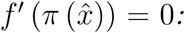: *in spite of a positive relatedness, uninvadability requires simple maximization of own fecundity. This holds true even if fecundity depends directly on the underlying trait, the standard assumption in the biology literature (see also Footnote 23), without being a function of some material payoff. It is this observation which is known as Taylor’s cancellation result, noticed initially in agent-based simulations by Wilson, Pollock, and Dugatkin (1992), proven formally by Taylor (1992a) for the island model, and then shown to hold for arbitrary migration patterns between groups (e.g., Taylor, 1992b, Rousset, 2004, and Ohtsuki, 2012). To see that it is in line with our result that κ* (*x*) = 0 *for s*_0_ = 0, *note that since f is strictly increasing in π*, 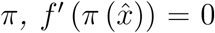 *in turn implies that* 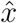 *maximizes π, i.e.*, 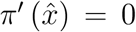. *Finally, we note that the same expression as that in the right hand side of eq. (29) was first obtained by Taylor and Irwin (2000, their eq. A.10), as a marginal cost to benefit ratio at the level of fecundity (see also Akçay and van Cleve, 2012). There is by now an extensive theoretical literature seeking to delineate how the assumptions pertaining to demography, life-history, the environment, and the modes of transmission, tip the balance in favor of pro- or anti-sociality at the survival or fecundity level (see, e.g., Eshel, 1972, Aoki, 1982, Gardner and West, 2006, Johnstone and Cant, 2008, Lehmann, Foster, and Feldman, 2008, Lion and Gandon, 2010, Bao and Wild, 2012, and Micheletti, Ruxton, and Gardner, 2017, for a some representative case studies, and Lehmann and Rousset, 2010, for a review).*

### 4.2 Scenario B: Guns

Take the biological scenario A with non-overlapping generations (set *s* (*π*) = 0 for all *π*), and augment it by introducing wars between groups. Following play of the material game in a demographic time period, but before reproduction, death of the adults, and migration by the offspring, islands are randomly engaged in pairwise wars, under exogenous uniform random matching. In each war, one island wins and the other loses. All individuals in the losing island thus die before they reproduce; the winning island takes over all reproductive resources of the other island and thus doubles its members’ fecundity. Technically, the double-sized pool of offspring of the winning island will split in two halves, one for each of the two islands, that they will treat as their “home” island. Let 0 ≤ *ρ* ≤ 1 denote the probability that any given island is drawn into war, the *war risk*, and let *g* (***π***, *π*^*^) denote the conditional probability that an island with material payoff profile *π* ∈ ℝ^*n*^ wins a war when the average payoff in the rest of the population is *π*^*^, conditional on being drawn into war. Here *g* is assumed to be increasing and permutation invariant with respect to the material payoffs earned by the inhabitants of the island in question. In other words, for *π*^*^ fixed, *g* has the properties of standard welfare functions. In this scenario the fitness function is

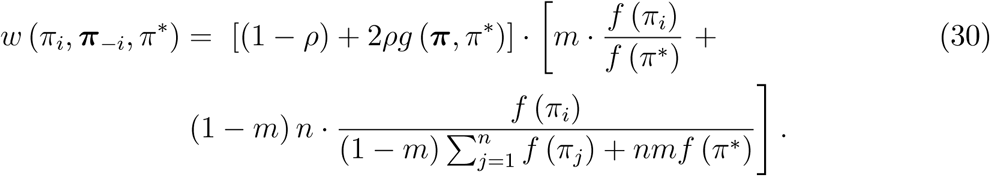

The difference with the baseline scenario is the first factor, which contains two terms: the probability that the individual’s island will not go to war (1 − *ρ*), and the probability that the island will go to war and win times two (2*ρg* (***π***, *π*^*^)), where the factor two comes from the assumption that a winning island doubles its fecundity and spreads its offspring uniformly over the two islands it now possesses. To see why the second factor is the same as the right-hand side of (26), note that migrants who arrive at any island, irrespective of whether this island has been involved in war or not, come with probability 1 − *ρ* from an island that was not in war, and (recalling that the average probability of winning a war is 1/2) with probability *ρ/*2 from an island that won a war. Moreover, victorious islands send out twice as many migrants as islands that did not go to war. Hence, the expected number of migrants who compete for the breeding spots in any given island is *m* (1 − *ρ* + 2*ρ/*2) · *f* (*π*^*^) = *mf* (*π*^*^), the same as in the absence of wars.

The coefficient of relatedness turns out to coincide with that in the preceding scenario (for *s*_0_ = 0). This is because the only event in which a randomly drawn individual can belong to the same local lineage as a randomly drawn neighbor, is when both belong to an island which did not lose a war, and both stayed in their natal island. Since the risk of losing a war applies to the whole island, while the migration probability applies to the individual, only the latter matters for relatedness. The coefficient of fitness interdependence equals

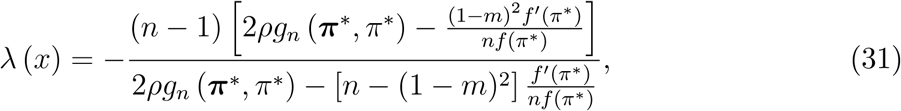

where *g*_*n*_ denotes the partial derivative of *g* with respect to the *n*-th argument (since *g* is evaluated in a homogenous population here, and since *g* is invariant under permutation of the *n* first arguments, *g*_*n*_ simply captures the marginal effect of an increase in the material payoff of any island member on the probability of winning a war). While the expression is involved, it can readily be seen (by considering a scenario in which *f′* (*π*^*^) = 0, for instance) that the effect of material payoffs on the strength in wars can make *λ* (*x*) negative, while in the scenario without wars studied above, it is always positive. In other words, conflicts between groups reduces spite, and may even reduce it so much that it turns into altruism, i.e., a positive weight is attached to the neighbors’ material payoffs. Indeed, by substituting (27) (for *s*_0_ = 0) and (31) into (22), we obtain:

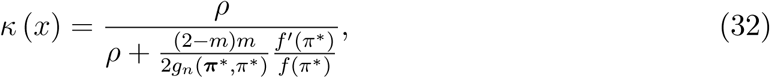

which is increasing in the marginal effect *g*_*n*_ on the probability of winning wars.

We next turn to weak selection in order to obtain more explicit results on the effects of wars on fitness interdependence and scaled relatedness. Recalling the notation under weak selection (see (16)), let each individual’s fecundity be exponentially increasing in the individual’s material payoff,

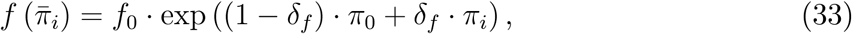

where *f*_0_ > 0 is baseline fecundity and *δ*_*f*_ > 0 represents the intensity of selection with respect to fecundity. Furthermore, assume that the probability of winning a war depends on the two islands’ aggregate material payoffs according to

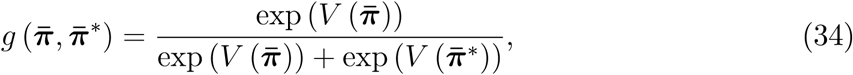

where 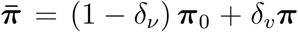 and 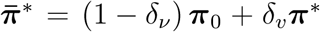, and *V* : ℝ^*n*^ → ℝ is a strictly increasing symmetric function (like any standard welfare function). Its values 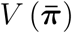 and 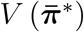 represent the “strengths” of the two islands. This is a logistic version of the Tullock contest function (Tullock, 1980), see Skaperdas (1996). It spans a continuum of cases, from all islands having the same chance to win any war, if the intensity of selection with respect to wars be nil, to the case in which the materially wealthiest island is almost sure to win any war (is the intensity of selection is infinitely large). Letting *δ*_*f*_ = *σ*_*f*_ · *δ* in equation (33) and *δ*_*v*_ = *σ*_*v*_ *δ*, for non-negative parameters *σ*_*f*_ ≥ 0, *σ*_*v*_ ≥ 0, and *δ* > 0, we can let both sensitivity parameters tend to zero at proportional rates by focusing on the limit as *δ* → 0. Below, however, we let *σ*_*v*_ = *σ*_*f*_, and thus write *δ* for *δ*_*v*_.

Many scenarios can be imagined, of which we consider two. First, if an island’s strength is proportional to its total material payoff, i.e., if 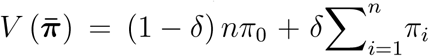, then fitness interdependence takes the following form (see the appendix):

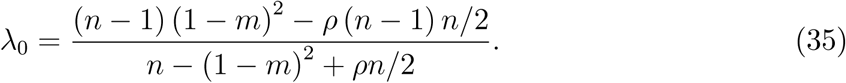

This changes sign when the risk of war is *ρ*^*^ = 2 (1 − *m*)^2^ */n*; it is positive at lower risks of war and negative at higher risk levels for war. Since in the baseline scenario with non-overlapping generations uninvadability under weak selection requires individuals to be selfish on balance (see Section 3.2), the reduction in fitness interdependence that the war risk entails, leads to pro-sociality on balance; indeed, for any *ρ* > 0 we obtain *κ*_0_ > 0:

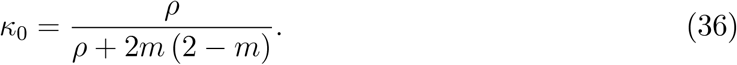

Moreover, the threat of war (*ρ* > 0) nourishes pro-sociality: *κ*_0_ is increasing in the risk of war, *ρ*, and is independent of group size, *n*.^24^ Figure 2 shows *κ*_0_ as a function of the migration rate *m*, for war risk *ρ* = 0 (the pink curve), *ρ* = 0.4 (the orange curve), and *ρ* = 0.8 (the blue curve).^25^

**Figure 2:**
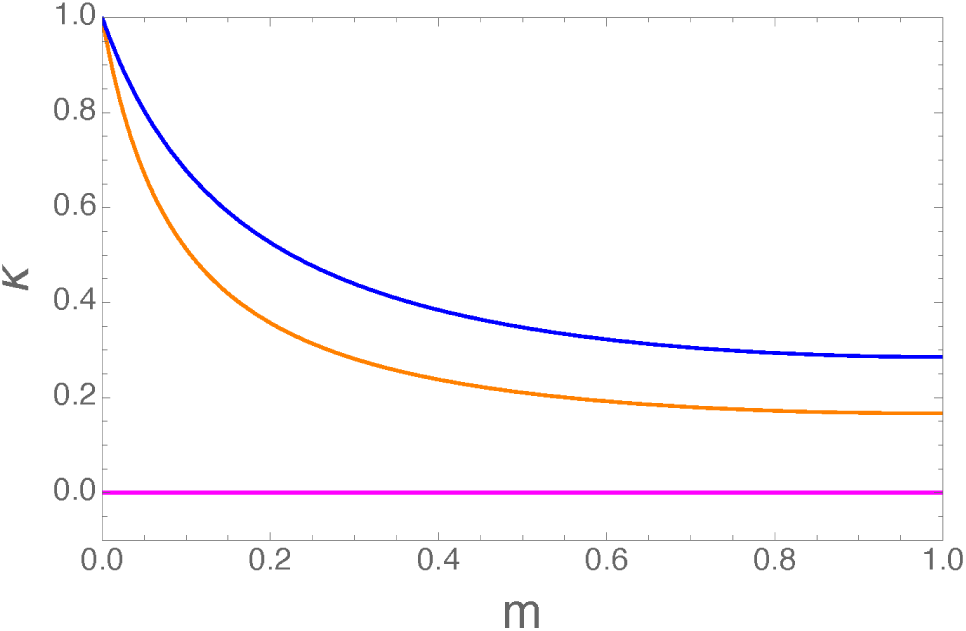
The value of *κ*_0_ as a function of the migration rate *m*.

Second, while it is arguably a natural benchmark case to assume that the probability of winning a war depends on the group’s total material payoff, sometimes the success or failure in conflicts depends on the strongest or the weakest member of one’s group.^26^ A general case, that allows for intermediate cases between dependence on the group’s total material payoff and its minimal payoff, is obtained by using a CES-functional form. Let

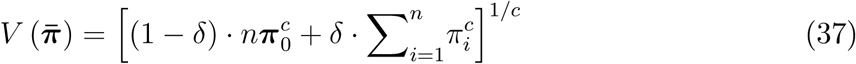

for some *c* ≠ 0. For *c* = 1 we obtain the previous case, and as 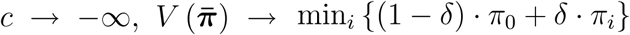 (Leontieff production function). Hence, when *c* is negative and large in absolute terms, an increase in the poorest group member’s material payoff will increase the winning probability, and hence have a positive effect on others’ fitness. This suggests a Rawlsian, rather than Benthamite concern for other group members’ material well-being. Individuals with medium or high material payoffs may then behave as if they had a particular concern for individuals with low payoff.

### 4.3 Scenario C: Culture

Suppose now that types are carried over from one generation to the next by cultural transmission. In every demographic time period, each adult dies and is replaced by exactly one child, who searches for a type to emulate, from its deceased (single) parent, another adult in its island, or an adult in another island. With probability *s* (*π*_*i*_) ∈ [0, 1], the loyalty of *i*’s child, the (unique) child of individual *i*, emulates its parent’s type. With probability 1 − *m* a non-loyal child searches for a type to emulate among the (now dead) grown-ups in its natal island (including its own parent). With the complementary probability, *m* > 0, such a child draws a sample of *n* grown-ups from the population at large, and emulates the type of one of them. The probability that an adult on any island is chosen as role model, when compared to others in her island (by a non-loyal child), depends on her type’s attractiveness relative to the attractiveness of the other grown-ups’ types in her island. Likewise, the probability that a child who searches outside its native island will pick a certain island, when looking for a “role model”, is assumed to be proportional to the island’s relative attractiveness in the world at large. Fitness *w* (*π*_*i*_, ***π***_−*i*_, *π*^*^) is then the expected number of children who emulate their type from an individual with material payoff *π*_*i*_ when the other island members earn the material payoff vector ***π***_−*i*_, and individuals in all other islands earn material payoff *π*^*^:

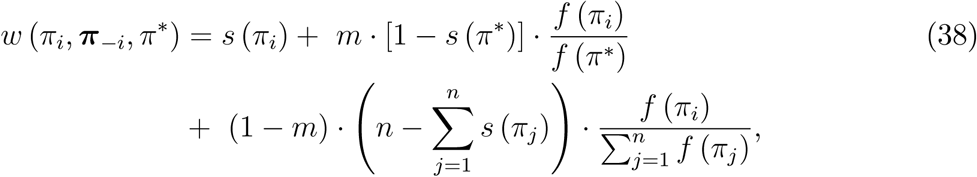

where, for any individual *j* in *i*’s island, *f* (*π*_*j*_) > 0 is the attractiveness of the type used by *j*. The first term in (38) is the probability that *i*’s child loyally emulates its parent’s type, without comparison with other adults’ types.^27^ The second term concerns the event that children from other islands emulate their type from one of the parents on *i*’s island. Written more explicitly, this term can be spelled out as

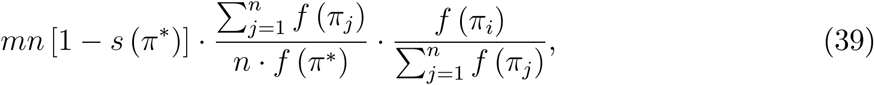

where the first factor is the expected number of children who search outside their native islands, the second factor is the probability for each such child to decide for *i*’s island, and the third is the conditional probability that it will then choose *i* as role model. The third term concerns the event that some or all the children in *i*’s island emulate their type from one among the parents on the island. The product of the first two factors in this term is the expected number of such children and the third factor is the probability, for each such child, that it will choose to imitate individual *i*. Note that, comparing this scenario to the biological scenario with overlapping generations, loyalty plays a similar role to survival, and attractiveness to fecundity. Moreover, the *cultural import propensity m* plays a similar role to migration. (These observations motivated the notation.)

In this scenario,

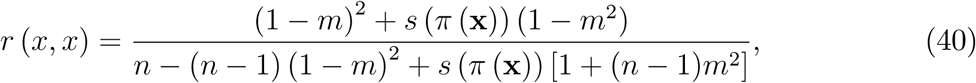

where x = (*x*, …, *x*) ∈ *X*^*n*^, and

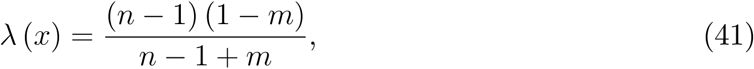

which leads to

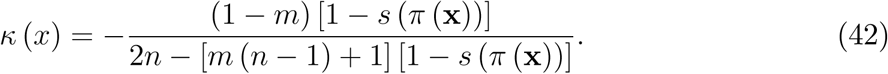

Comparison with the biological scenario with overlapping generations reveals that the coefficients of relatedness are the same, but that for any *m* < 1 the coefficient of fitness inter-dependence is larger under cultural transmission. The enhanced competitiveness is strong enough to lead to anti-sociality, since *κ* (*x*) < 0 obtains if and only if (1 − *m*) [1 − *s* (*π* (x))] < (2 − *m* [1 − *s* (*π* (x))]) · *n*, an inequality which holds for all parameter values^28^. In this example, cultural transmission thus leads to anti-sociality, and anti-sociality is stronger at low values of *m*. This is because a low cultural import rate enhances fitness interdependence. Note that although genetic and cultural transmission here lead to opposite predictions regarding sociality, one qualitative similarity that appears is that like survival under genetic transmission, loyalty under cultural transmission has a positive effect on sociality, *κ* (*x*). We also note that the negative pro-sociality vanishes as groups tend to become infinitely large: *κ* (*x*) → 0 as *n* → ∞.

To illustrate this, Figure 3 shows that *κ* (*x*) is strictly negative for all *m* < 1, for different loyalty rates and different island sizes: for *s*_0_ = 0 and *n* = 2 (the pink curve), *s*_0_ = 0.4 and *n* = 2 (the orange curve), *s*_0_ = 0.8 and *n* = 2 (the blue curve), *s*_0_ = 0 and *n* = 10 (the pink dashed curve), *s*_0_ = 0.4 and *n* = 10 (the orange dashed curve), *s*_0_ = 0.8 and *n* = 10 (the blue dashed curve).

**Figure 3:**
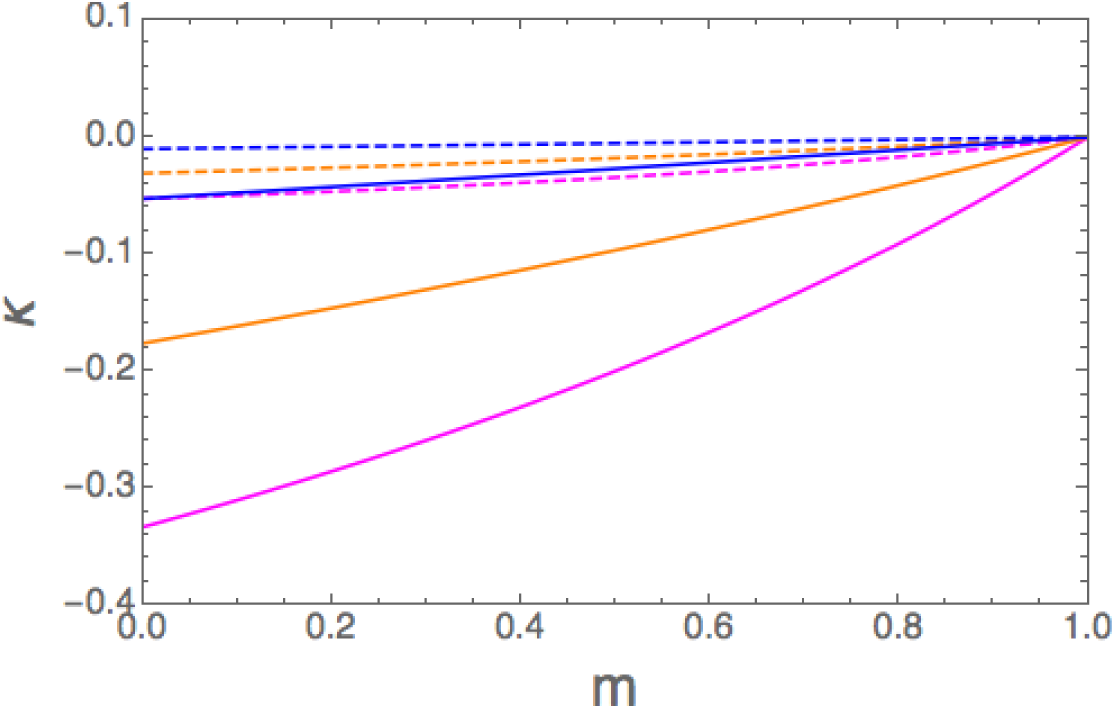
The value of *κ* (*x*) as a function of the cultural import rate *m*, for different degrees of background loyalty of offspring towards parents.

## 5 Conclusion

By combining non-cooperative game theory and evolutionary biology, we have derived several novel insights on the evolutionary viability of preferences in social interactions. In particular, our model enables analysis of how the tendency of individuals to interact in fairly small groups, between which there is limited migration, and between which there may be conflicts, affects such preferences. A key strength of the model is that it makes a distinction between material payoffs, which typically is the level at which data analysis by economists is conducted, and individual fitness. Our results clearly show that the qualitative nature of evolutionarily viable preferences is typically different at the material payoff than at the individual fitness level. Furthermore, our results provide an evolutionary justification for preferences as drivers of choice, by connecting stability at the strategy level with equilibrium behavior under certain preferences. Our results thus address a criticism of the literature on preference evolution, according to which it conflates revealed preferences with preferences that drive choice, see, e.g. Newton (2018).

The cognitive assumption we make is that individuals understand what interaction is at hand, but they need not know the material payoffs to others or the preferences of others. Moreover, our formalization allows for the possibility that in fact there are (finitely) many interactions going on simultaneously, or that are randomly selected, and even that each interaction involves only a subset of the inhabitants in an island. What is required is symmetry in the sense that all individuals face the same probabilities of being involved in any one of the interactions and that the interaction at hand is aggregative and symmetric.

However, if individuals also understand the mapping from strategies to material payoffs, a remarkable result emerges from our analysis. Under weak selection the nature of the derived preferences are independent of the nature of the strategic interaction within islands. This is because the matching profiles then depend only on the population structure, without any reference to material payoffs. Hence, the utility function *v*^0^ (see (18)) would remain uninvadable even if the mapping from material payoffs to fitness and/or the mapping from strategies to material payoffs were to change over time, as long as these changes do not affect the matching probabilities, and as long as individuals understand the mapping from strategies to material payoffs and adjust the material payoff terms in *v*^0^ accordingly.^29^ Such robustness, however, presumes that Nash equilibrium play under the adjusted *v*^0^ would be reached. Furthermore, the utility function at the fitness level would generally not remain uninvadable. Given that the aforementioned mappings have certainly changed over the course of human history, future research should lift the assumption of time-invariant mappings.

While our model is general in the sense that we allow for essentially any type of interactions within groups, it also has several limitations. Perhaps the strongest is that we only analyze type-homogeneous play and homogeneous populations subject to a single mutant. More realistic models, with heterogeneous individuals, heterogeneous islands and resident populations with multiple types are called for. Our hope is that the model proposed here can be fruitfully used to this end.

## 6 Appendix

### 6.1 Fitness and randomness

We here give details about how we justify the expression of individual direct fitness *w* (*π*_*i*_, ***π***_−*i*_, *π*^*^) in our model. First, we note that, generically in the infinite island model, lineage fitness ofa mutant trait *τ* in a resident population with trait *θ* is defined as

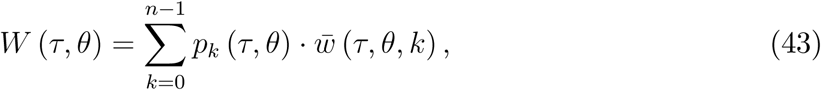

where 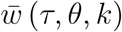 is the expected number of settled offspring in the next demographic time period that descend from a given adult mutant with trait *τ* in a group with exactly *k* other mutants, and thus *n* − *k* − 1 individuals with trait *θ*. More formally, 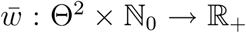 is defined as the expectation of the random number *W*_*τ,θ,k*_ ∈ ℕ_0_ of settled offspring descending from the given mutant (including herself through survival), conditional on the event that in the parental demographic time period her island is in *state s* = (*τ, θ, k*), that is, with *k* other mutants (with trait *τ*) and the other *n* − *k* − 1 individuals with trait *θ*. The stochasticity in the random variable *W*_*τ,θ,k*_ is due to *within-generation variability*.

Let **Π** = (Π_1_, Π_2_, …, Π_*n*_, Π^*^) be the *random payoff vector* on an island, where Π_*i*_ is the random material payoff obtained by individual *i* = 1, …, *n* at the end of phase one of a demographic time period, and Π^*^ is the random payoff earned by a representative individual in an island where all individuals have trait *θ*. According to our decomposition of a demographic time period into two phases (see section 2.1):

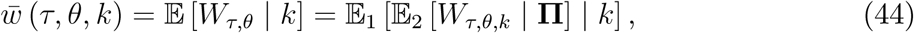

where 𝔼_1_ is the expectation over all stochastic events occurring during phase 1 of the demographic time period (potential randomness in the actions taken by individuals, and hence in payoffs obtained), while 𝔼_2_ is the expectation over all stochastic events occurring during phase 2 of the demographic time period (randomness in reproduction, survival, and/or sampling among competing offspring).

We note that three sources of within-generation variability can be distinguished in our model: (i) within-island *trait variability* (randomness in the number of other mutants), (ii) within-island *interaction and payoff variability* (for given number of mutants, randomness in the payoff vector), (iii) within-individual variability (for given number of mutants and payoffs, randomness in survival and number of surviving offspring). Hence, equation (43) can be viewed as a three-level iterated expectation:

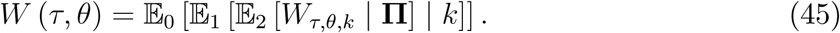

This is the grand expectation of the random number *W*_*τ,θ*_ of settled offspring descending from a mutant randomly sampled from the local lineage of the initial mutant, sampled during the random time interval until the first extinction of the local lineage, a time interval that is finite with probability one.

We are now in a position to introduce the continuously differentiable *individual fitness function w* : ℝ^*n*+1^ → ℝ that maps realized material payoff vectors to the expected number of offspring, conditional to the island state *s* = (*τ, θ, k*):

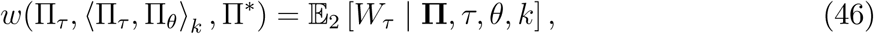

where ⟨Π_*τ*_, Π_*θ*_⟩_*k*_ is the random vector of the island neighbors’ payoffs, when *k* neighbors (of the given mutant) are mutants and the others are residents. We note that in a homogeneous population, that is, where all individuals carry the same trait, irrespective what that trait is, and all individuals use the same strategy, the random payoffs are identically and independently distributed, and hence, for 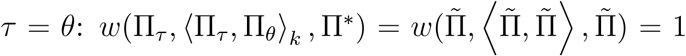, due to the constancy of the population in our model. Hence,

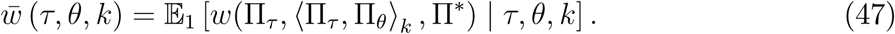

### 6.2 Functions and randomness

So far, we imposed no restrictions on the effect of within-generation uncertainty. A key assumption we make in the analysis in the main text is that

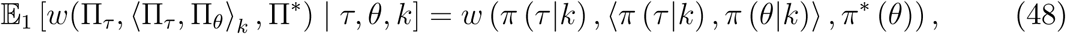

where *π* (*τ* |*k*) = 𝔼_1_ [Π_*τ*_|*τ, θ, k*], *π* (*θ* |*k*) = 𝔼_1_ [Π_*θ*_|*τ, θ, k*], *π*^*^ = 𝔼_1_ [Π_*θ*_|*θ, θ, k*]. Hence, we replace the expectation of a function by the function of the expectation for uncertainty in phase 1 (type (ii) uncertainty above), which is a substantial assumption, except when all functions are affine.

When the game under consideration is but one source for individuals’ fitness and there is variance in payoff, equation (48) is less restrictive under weak selection than may first be thought. To see this, suppose that the total random payoff to an individual *i* (who may have trait *θ* or *τ*) is the convex combination of two random variables, one exogenous random background payoff (from other interactions, say), Π_*b*_, and the random payoff 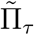 from the material game under consideration in our model:

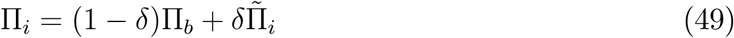

where *δ* ∈ (0, 1) is small. Then, by way of a Taylor expansion with respect to *δ* at *δ* = 0, and using the zero-sum property of effects on individuals’ fitnesses, we get (with subscripts on the function *w* denoting partial derivatives):

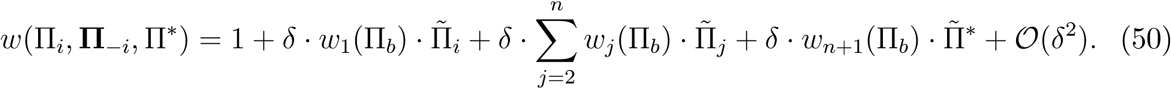

Suppose further that the random baseline payoff Π_*b*_ is statistically independent from that of the specific game, which for an mutant individual *i* denote 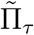. Then

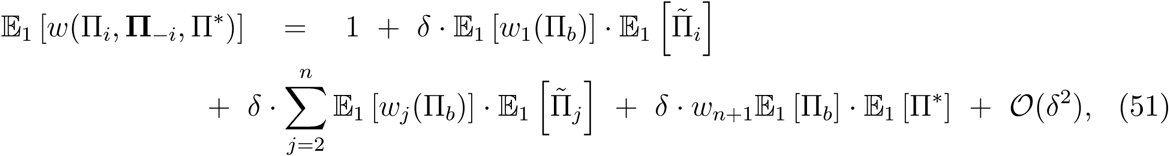

which leads to the same results as obtained by eq. (68) here below, but with partial derivatives *w*_*j*_(Π_*b*_) replaced by their expectation.

### 6.3 Proof of Proposition 1

We show first that 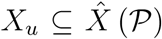 is a sufficient condition for *u* to be uninvadable. Suppose that 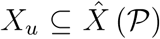. Then for each 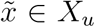, (8) is satisfied for any strategy *y* ∈ *X* played by mutants. In other words, there exists no *v* ∈ ℱ for which some (*x, y*) *B*_*NE*_ (*u, v*) does not satisfy the inequality in (10). Hence, the condition (10) for *u* to be uninvadable in Θ = ℱ is satisfied.

We now show that 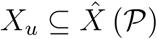 is a necessary condition for *u* is uninvadable. Suppose to the contrary that *u* is uninvadable and that there exists some 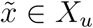 such that 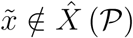. Then there exists some 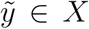 for which the inequality in (10) is not satisfied for 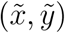. Consider the mutant utility function 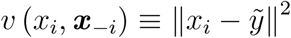; it induces mutants to play the strategy 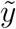 whichever strategy the residents play. Hence, there exists (*x, y*) ∈ *B*_*NE*_ (*u, v*) for which (10) is not satisfied. Since *v* ∈ ℱ, this means that *u* is invadable in Θ = ℱ.

### 6.4 Proof of Proposition 2

Consider some uninvadable strategy 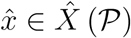. Then

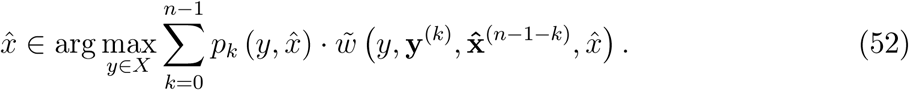

Suppose now that 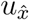 is the resident utility function. To see that 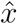 is then a resident strategy, note that given that an individual *i*’s opponents in the group play 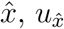 writes:

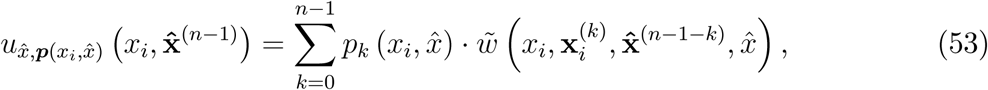

so that 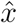 is a resident strategy iff

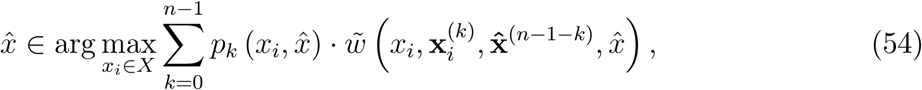

which is true (to see this, compare this expression to (52)).

However, the fact that 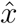 is the unique strategy satisfying (52) does not preclude existence of other resident strategies under 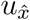. Indeed, consider some strategy 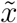. This is a resident strategy if

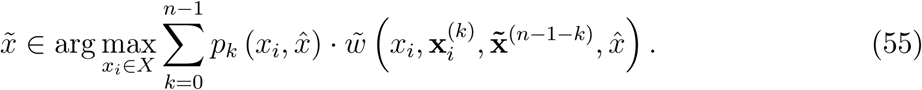

Lastly, if 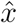 is the unique resident strategy under 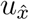, then the set of resident strategies under 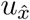 is a subset of 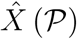. This together with Proposition 1 implies that 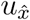 is uninvadable in ℱ.

### 6.5 Proof of Proposition 3

For *x* to be uninvadable it must be that, given *x, y* = *x* is a local maximum of

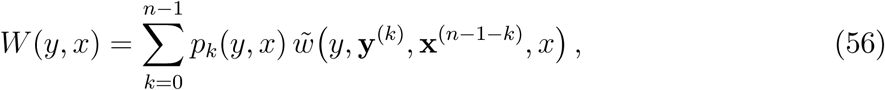

where **y**^*(k)*^ is the *k*-dimensional vector whose components all equal *y*, and **x**^*(n*−1−*k)*^ is the (*n* − 1 − *k*)-dimensional vector whose components all equal *x*, or 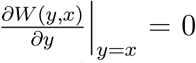. To evaluate this first-order condition, we follow the same calculations as in Lehmann, Alger, and Weibull (2015) Appendix B. In particular, writing 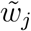 for the partial derivative of 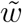 with respect to its *j*-th argument,

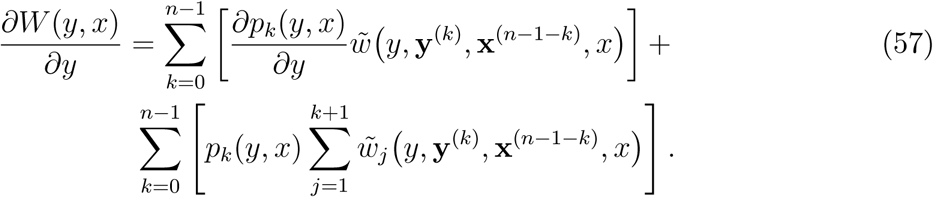

Noting that for 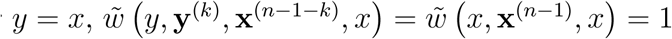, which is independent of *k* so that it can be factored out in the first term, and that

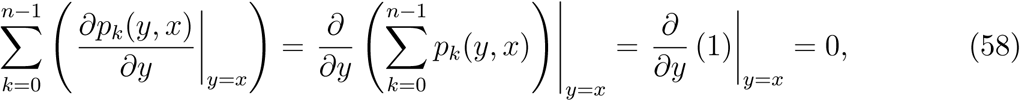

the expression simplifies to

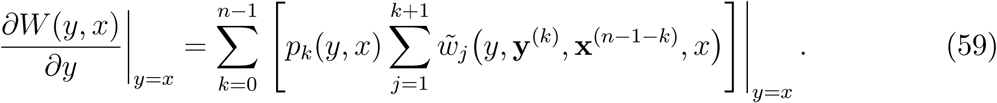

Permutation invariance further implies that for any 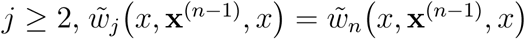 (it’s as if the individual whose marginal type change is under consideration were systematically labeled to appear as the last component in the vector x^*(n*−1)^). Noticing also that 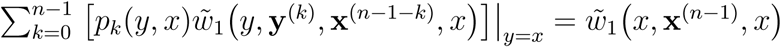, we can write:

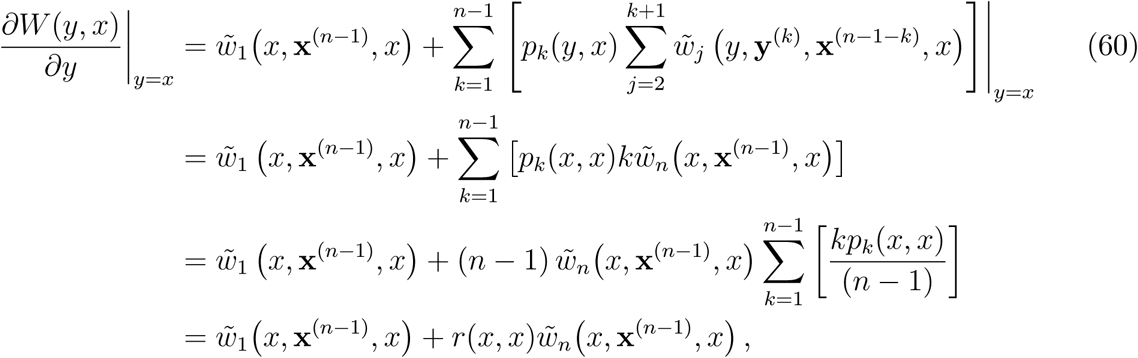

which owing to permutation invariance can also be written

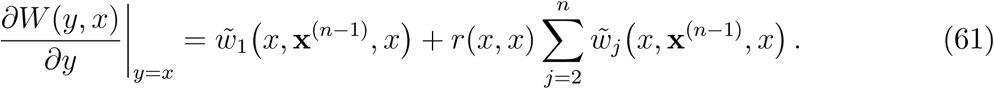

### 6.6 Proof of Proposition 4

The proof begins by deriving a lemma under strategy evolution, which is a generalization of Appendix B of Lehmann, Alger, and Weibull (2015), and will be a stepping stone towards the result on preference evolution stated in the proposition. For this purpose, we define the *lineage payoff-advantage* of a mutant strategy *y* ∈ *X* in a population of residents using strategy *x* ∈ *X* as

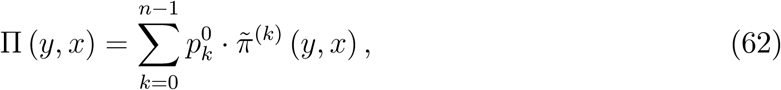

where 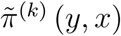 is the mutant’s *payoff advantage* when there are *k* other mutants in her or his island, defined by

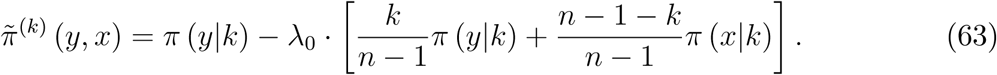

The first term in (63) is the payoff of a descendant of the initial mutant who finds herself in an island with *k* other such descendants. The term in square brackets is the average material payoff earned by the other members in the island.

#### Lemma 1

*A strategy* 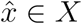 *is uninvadable under weak selection if and only if*

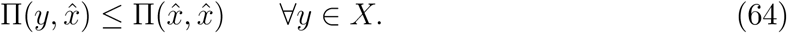

*Moreover*, 1 − *n* ≤ *λ*_0_ ≤ 1.

**Proof of Lemma 1:** Let *w* : ℝ^*n*+1^ → ℝ be any continuously differentiable fitness function, let *b* ∈ ℝ, and let b denote the vector in ℝ^*n*+1^ that has all components equal to *b* ∈. Then, by virtue of (86),

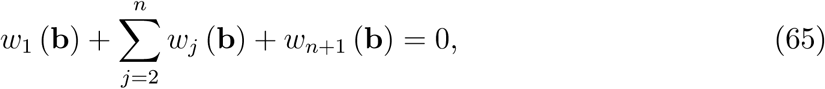

where an index *k* = 1, …, *n* +1 stands for the partial derivative of *w* with respect to its *k*-th argument.

Recalling the definition of 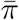 (see (16)), and omitting for notational simplicity the term (1 − *δ*) *π*_0_, for any given payoff vector (*π*_*i*_, ***π***_−*i*_, *π*^*^) ∈ ℝ^*n*+1^ a first-order Taylor expansion of *w* with respect to *δ* evaluated at *δ*_0_ writes

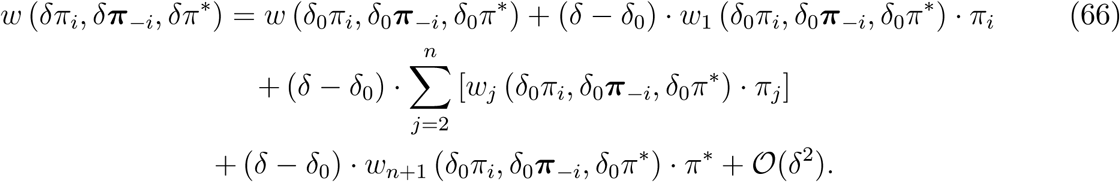

Evaluated at *δ*_0_ = 0, this expression writes

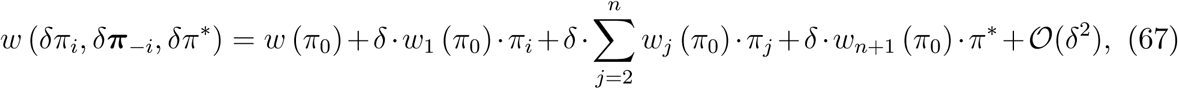

where *w* (*π*_0_) = 1, and *π*_0_ is the vector in ℝ^*n*+1^ whose components all equal *π*_0_. By permutation invariance of *w* (*π*_*i*_, ***π***_−*i*_, *π*\*) with respect to the components of ***π***_−*i*_, we may for each *j* = 2, …, *n* write *w*_*n*_ (*π*_0_) instead of *w*_*j*_ (*π*_0_). Letting *β* = *w*_1_ (*π*_0_) and *γ* = − (*n* − 1) *w*_*n*_ (*π*_0_), using (65), and rearranging terms, (67) can thus be written

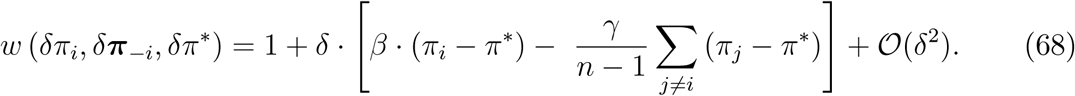

Letting

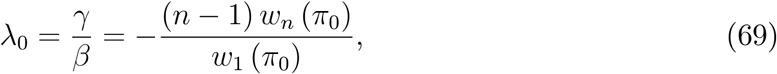

and factoring out *β* > 0 from (68), and simply omitting to write the factor *δ* in the fitness function, we conclude that for small *δ* > 0,

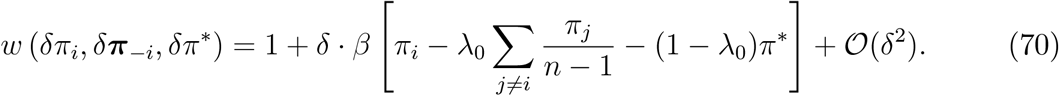

This shows that *λ*_0_ quantifies fitness interdependence among patch members (Lehmann, Alger, and Weibull, 2015; see also Frank, 1998, and Gardner and West, 2004, for a description, but without a formal derivation, of *λ*_0_).

The next step of the proof consists in obtaining an expression for local linage fitness under weak selection. Under weak selection the evolutionary process is what in biology is called *neutral* (Crow and Kimura, 1970, Ewens, 2004, Gillespie, 2004, and, for an explicit example, Rousset, 2004). Formally, this means that we can write

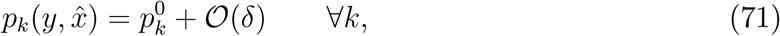

where 𝒪(*δ*) accounts for the deviation (relative to the neutral process) of the strategy-profile distribution induced by selection (i.e., *δ* > 0) that is at most of order *δ*, where 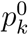 is strategy-independent. Second, recalling the definition of 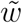 (see (7)) and letting 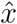 denote the resident strategy, (70) can be written

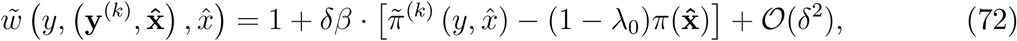

where 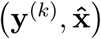 is the (*n* − 1)-dimensional vector with *k* components equal to *y* and the remaining components equal to 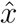, and (see equation (63))

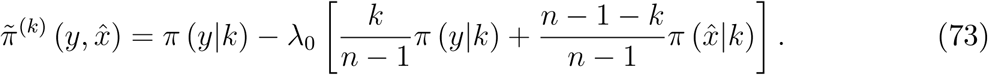

Using (71) and (72), local lineage fitness (see (2)) writes

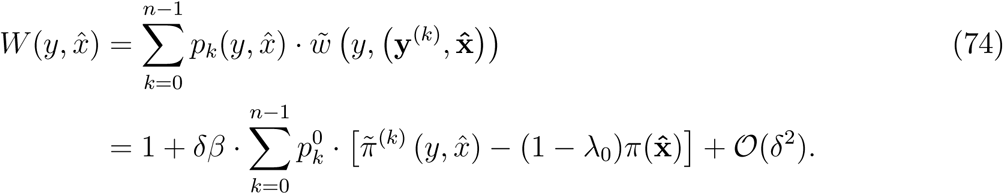

Recalling the definition of lineage payoff-advantage 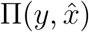 (see (62)),this can be written as

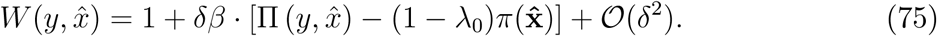

Neglecting higher order terms in *δ* in this equation, the condition for uninvadability 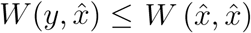 for all *y* ∈ *X*] under weak selection is equivalent to the condition 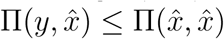 for all *y* ∈ *X*.

Finally, we determine the implications of Assumption [**M**] for the bounds on *λ*_0_ = − (*n* − 1) · *w*_*n*_ (*π*_0_) */w*_1_ (*π*_0_), focusing on the non-trivial case *w*_*n*_ (*π*_0_) ≠ 0. Part *(ii)* of the assumption implies − (*n* − 1) ≤ *λ*_0_. Moreover, recalling (65) we obtain *λ*_0_ ≤ 1, with strict inequality when either *w*_*n*+1_ (*π*_0_) < 0 or *w*_*n*+1_ (*π*_0_) = 0 and *n* > 2. **Q**.**E**.**D**.

We are now in a position to complete the proof of the proposition. To establish the first claim of the proposition, we note that Lemma 1 implies that 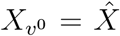. The second claim follows by noting that any utility function *u* ∈ ℱ for which some *x* ∈ *X*_*u*_ is not an element of 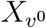, is invadable.

### 6.7 Approximation of the neutral distribution

Standard populations genetics results (see e.g., Lessard, 2007, and references therein) show that the neutral distribution of types in an island model with constant group size, and with population share of mutants *ε* > 0, is well approximated by way of the hypergeometric distribution

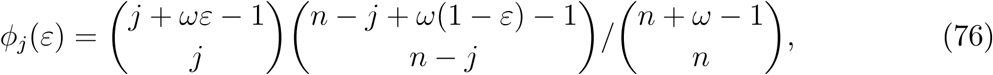

where *ϕ*_*j*_(*ε*) is the probability that there are *j* = 0, 1,…, *n* mutants in any given group, and *ω* = *r*_0_*/*(1 − *r*_0_) (see Lessard, 2007, equation (7)). Since 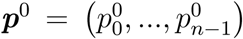 is the limit distribution when *ε* → 0 of the number of *other* mutants in a given mutant’s group, we have

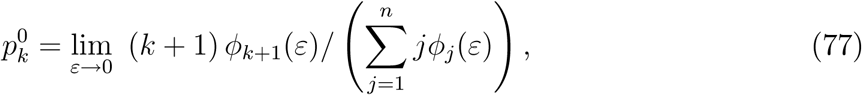

for *k* = 0, 1, …, *n* − 1. Upon rearrangements, this produces

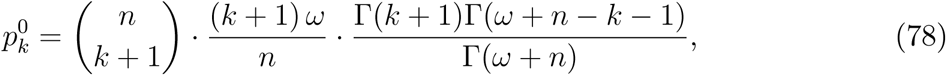

where Γ is the gamma function. This distribution depends only on group size *n* and pairwise relatedness *r*_0_.

Numerical comparison between this approximation for the above evolutionary scenarios (that can all be subsumed under the relatedness in (40)) and the exact distribution shows that the average total variation between the approximate and exact distributions is quite small. Sampling randomly 10 000 values of *s* and *m* when *n* = 5 gives an average total variation of 0.005, a variation that should diminish with *n*. It can also be shown that in the special case of a Moran process (*s* (*π*_*i*_) = 1*/n*) the approximation is in fact exact. (Indeed it can be verified that the expression in (78) then reduces to equation D.6 in Lehmann, Alger, and Weibull, 2015.)

### 6.8 Proof of Proposition 5

Recalling that

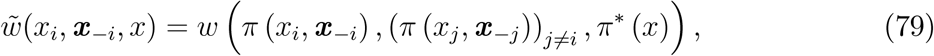

we obtain

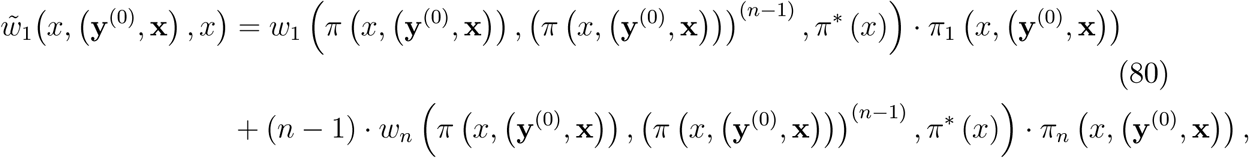

where (*π*(*x*,(**y**^(0)^ **x**)))^(*n*−1)^ denotes the (n − 1)-dimensional vector whose components all equal *π* (***x***, (**y**^(0)^, **x**)), and

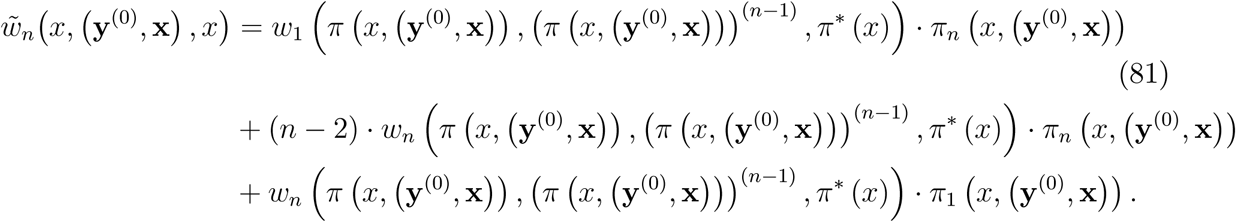

Substituting the last two equations into the last line of (60) produces

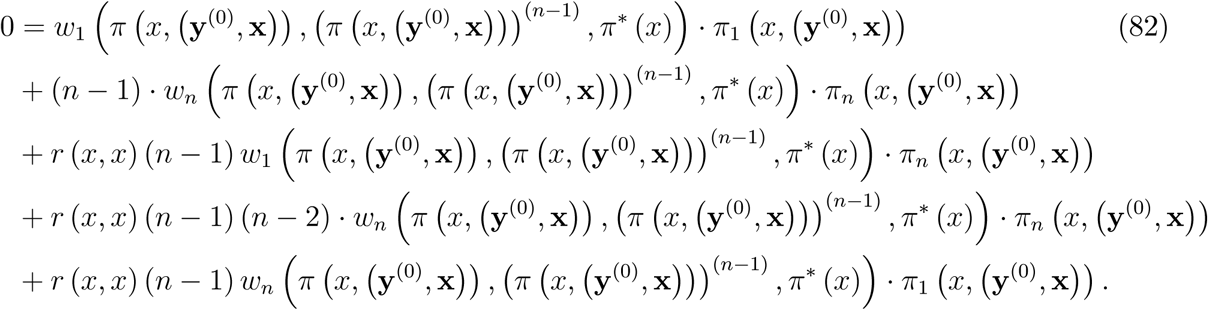

Noting that with the notation used in this proof, *λ* (*x*) writes

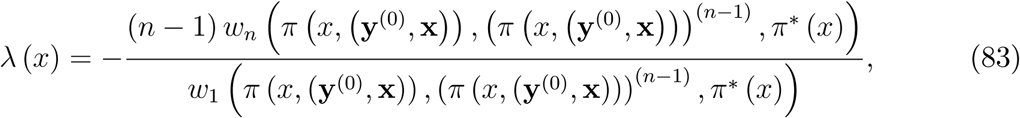

(82) can be written

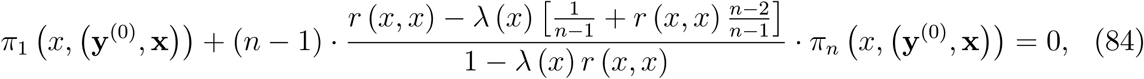

or

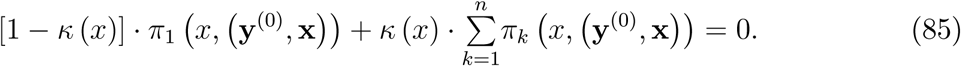

### 6.9 Proof of Proposition 6

To show that *κ* (*x*) ∈ [−1, 1], we begin by studying *λ* (*x*). Note that the terms that define *λ* (*x*) are partial derivatives evaluated in a homogenous population. Furthermore, since population size is constant in a homogenous population, each individual’s fitness would remain at 1 following a marginal change in the material payoff of all the individuals in the population. Formally:

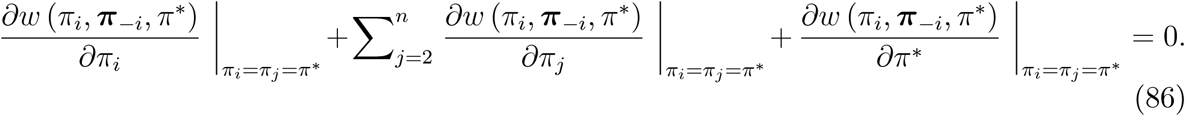

By permutation invariance, and using a more compact notation, this writes *w*_1_ (·)+(*n* − 1) *w*_*n*_ (·)+ *w*_*n+1*_ (·) = 0. Using this and the assumption *w*_1_ (·) > 0,

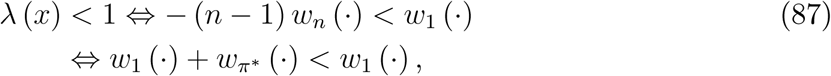

which is true by Assumption [**M**] *(iii)*.

Since *r* (*x, x*) ∈ [0, 1] for all *x* this implies that *λ* (*x*) *r* (*x, x*) < 1, and hence

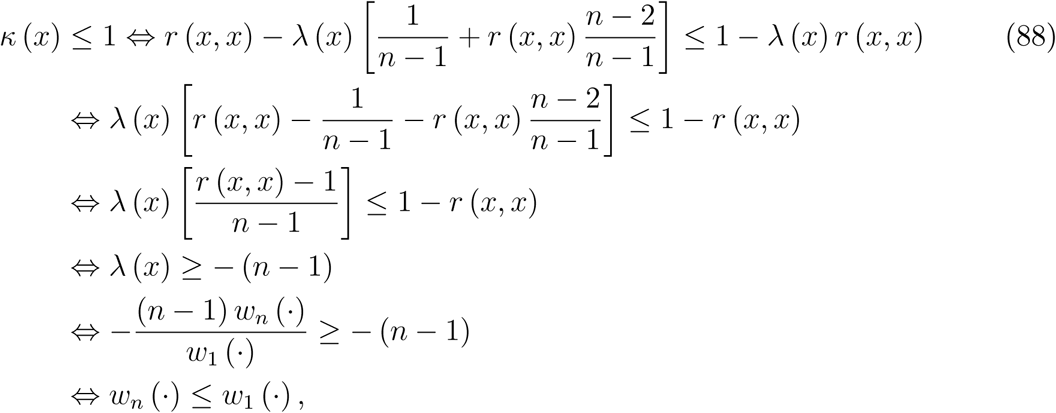

which is true by virtue of Assumption [**M**] *(ii)*.

We now show that *κ* (*x*) ≥ − 1. For any *λ* (*x*) < 1, *κ* (*x*) is increasing in *r* (*x, x*). Indeed, the partial derivative of the expression for *κ* (*x*) with respect to *r* (*x, x*) has the same sign as (in this expression *r* ≡ *r* (*x, x*) and *λ* ≡ *λ* (*x*))

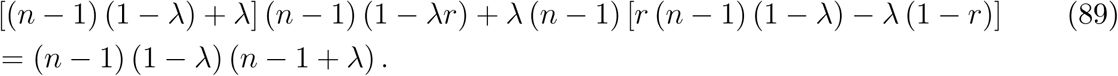

For the inequality *κ* (*x*) ≥ −1 to hold, it is thus sufficient that *κ* (*x*) ≥ −1 for *r* (*x, x*) = 0, a condition which reduces to

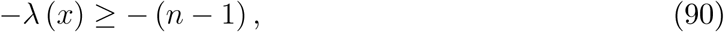

which is true for any *n* ≥ 2 since *λ* (*x*) < 1.

Finally,

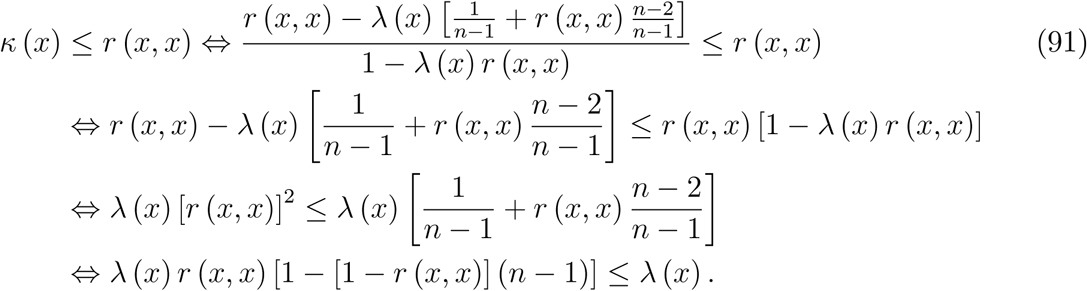

This inequality is true if and only if *λ* (*x*) ≥0 by virtue of the fact that for all *r* (*x, x*) ∈ [0, 1) we have *r* (*x, x*) [1 − [1 − *r* (*x, x*)] (*n* − 1)] ≤1. Likewise, it is clear that *κ* (*x*) > *r* (*x, x*) if and only if *λ* (*x*) < 0.

Finally, the last result stated in the proposition is implied by (20) together with Assumption [**M**] *(i)*.

### 6.10 Calculating the coefficients of fitness interdependence and pairwise relatedness

#### 6.10.1 Scenario A: Genes

To calculate *λ* (*x*) we begin by calculating the partial derivatives needed for this purpose. Here, from the individual fitness function (26):

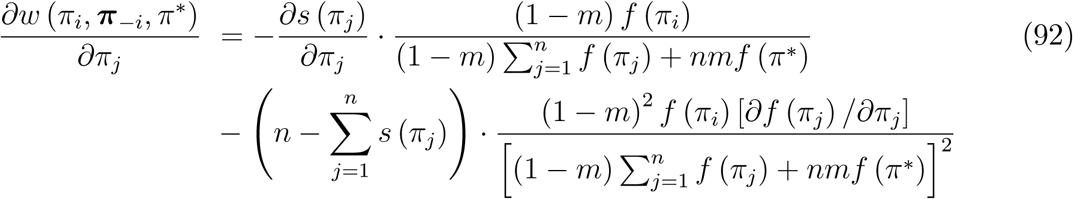

and

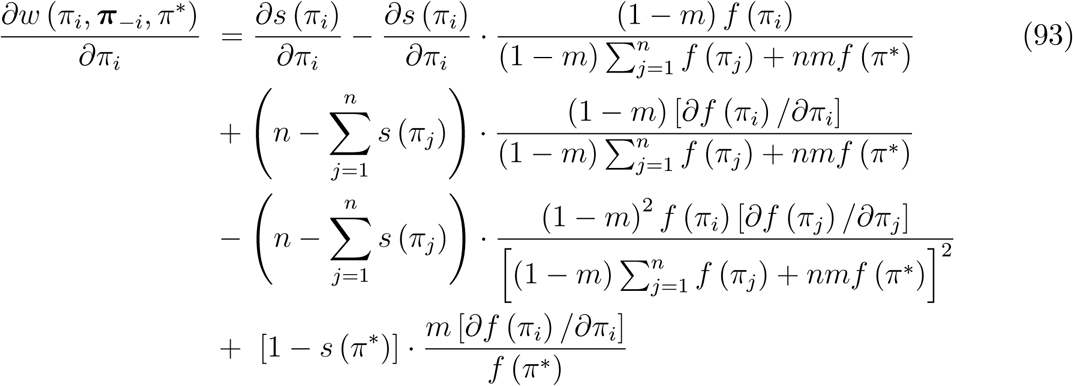

Writing *s*′ (*π*\*) for 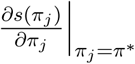 and *f*′ (*π*\*) for 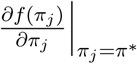, we obtain (upon simplification)

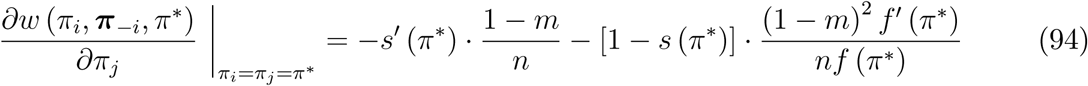

and

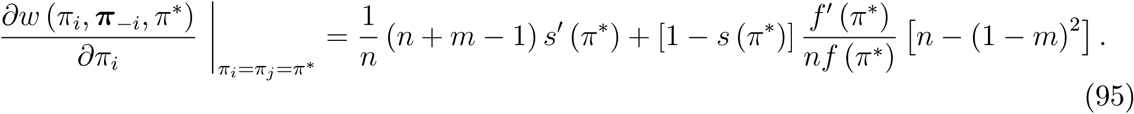

Upon simplification, we thus obtain

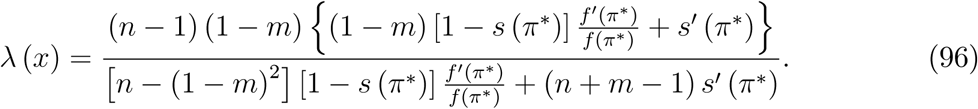

The expression in (28) obtains by setting *s*′ (*π*^*^) = 0.

To calculate *r* (*x, x*), one uses a recursion equation (this is a standard technique for calculating probabilities of identity by descent; see Nagylaki, 1992, and Rousset, 2004, for a background). In the scenario at hand, this equation writes

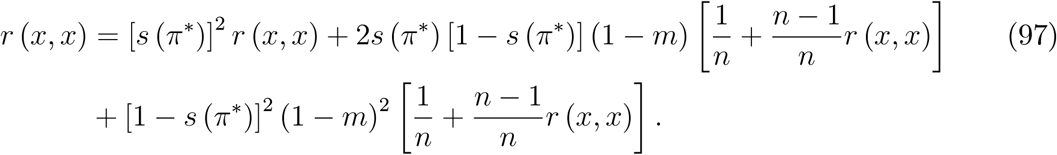

The left-hand side is the average probability that, in a monomorphic population in which all individuals play *x*, the neighbor of a randomly drawn member of a certain local lineage is also a member of this local lineage. The terms on the right-hand side details the events in which this happens. The first term on the right hand side corresponds to the event that both the individual at hand and the randomly drawn neighbor survived from the previous period. The second term on the right hand side corresponds to the two events in which either the individual at hand or the randomly drawn neighbor survived from the previous period while the other didn’t, and the one who didn’t survive from the previous period did not migrate in from another island. In this case, there is a probability 1*/n* that one is the offspring of the other, in which case they are both members of the same local lineage with certainty; with the complementary probability, they are not parent and child, in which case the probability that they are both members of the same local lineage equals *r* (*x, x*). The third term on the right hand side corresponds to the event in which neither the individual at hand nor the randomly drawn neighbor survived from the previous period, and neither of them migrated in from another island. In this case, there is a probability 1*/n* that they have the same parent, in which case they are both members of the same local lineage with certainty; with the complementary probability, they have different parents, in which case the probability that they are both members of the same local lineage equals *r* (*x, x*). Solving (97) for *r* (*x, x*) yields

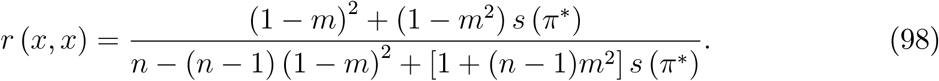

The expression in (27) obtains by replacing *s* (*π*^*^) by *s*_0_.

#### 6.10.2 Scenario B: Guns

In the biological scenario with wars, we obtain from the individual fitness function (30):

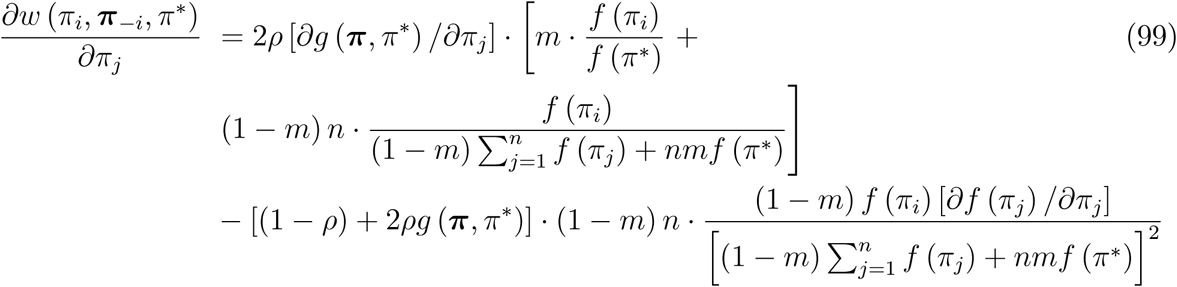

and

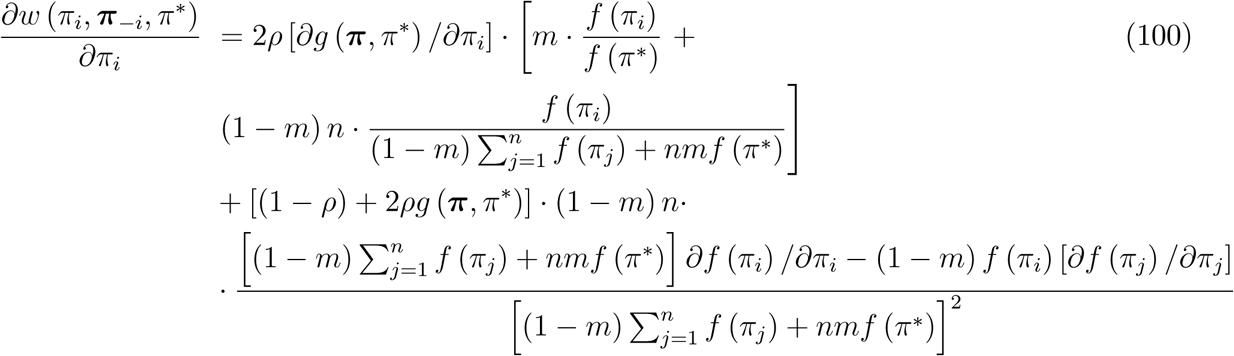

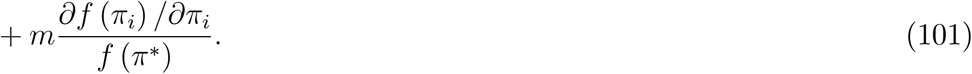

By permutation invariance of *g*, write *g*_*n*_ (***π***^*^, *π*^*^) for 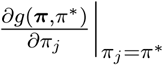 for all *j* = 1, …, *n*. Since, moreover, *g* (***π***, *π*^*^) = 1*/*2, upon simplification we obtain:

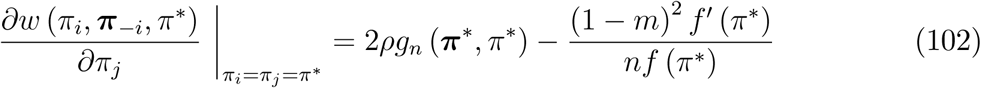

and

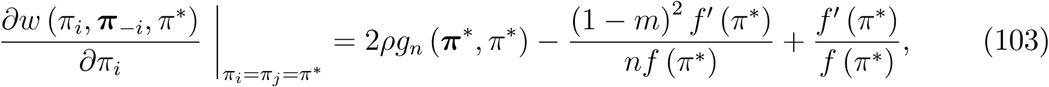

so that

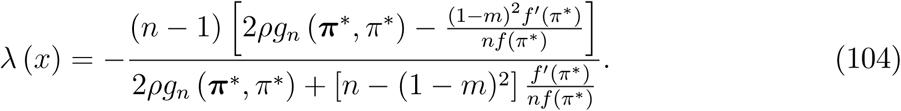

The recursion equation to calculate *r* (*x, x*) writes

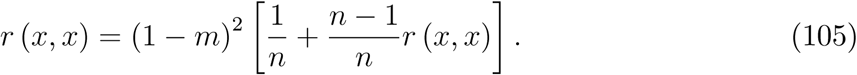

In this scenario, the only event in which a randomly drawn individual can belong to the same local lineage as a randomly drawn neighbor, is when both stayed in their natal island. In this case, there is a probability 1*/n* that they have the same parent, in which case they belong to the same local lineage with certainty; with the complementary probability, they have different parents, in which case the probability that they belong to the same local lineage is *r* (*x, x*). Solving for *r* (*x, x*) yields

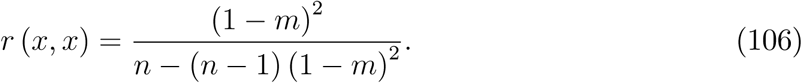

#### 6.10.3 Scenario B: Wars (weak selection)

Recall that under weak selection we write the individual fitness of individual *i* as 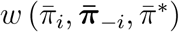, where 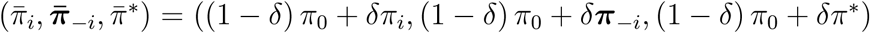, and *δ* ≥ 0 represents the intensity of selection (see (16)). From (69) in the proof of Proposition 5, we have

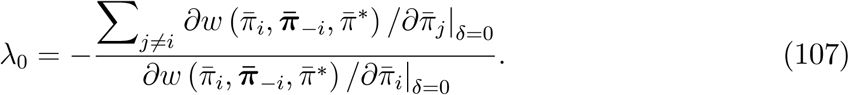

Since, for *δ* = 0, 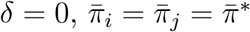, we obtain from (102) and (103) that

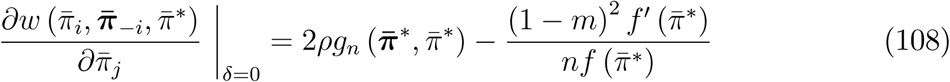

and

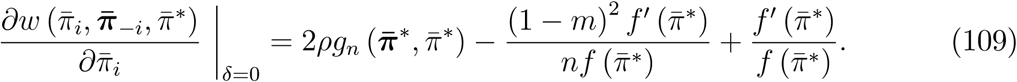

With the expressions for *f* and *g* given in (33) and (34), and the assumption that 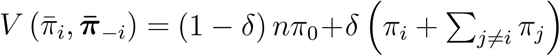 (note that we assume that the intensity of selection is the same for fecundity and for the probability of winning wars; one can also allow for different selection intensities), we have:

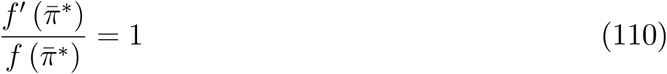

and

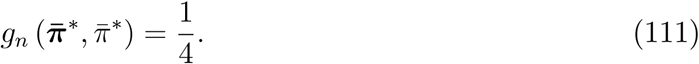

Hence, we get

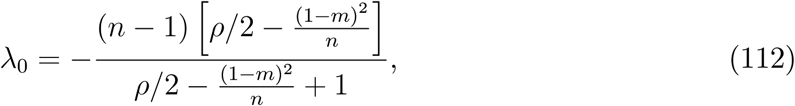

which upon simplification gives the expression in (35). It can then be verified that, together with the fact that *r*_0_ is given by (106), this gives the expression for *κ*_0_ in (36).

#### 6.10.4 Scenario C: Culture

In the cultural scenario, we have from (38):

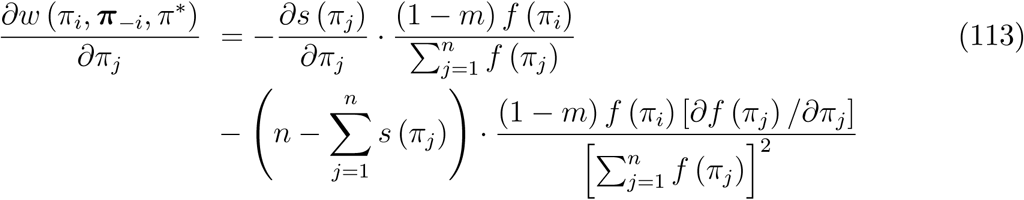

and

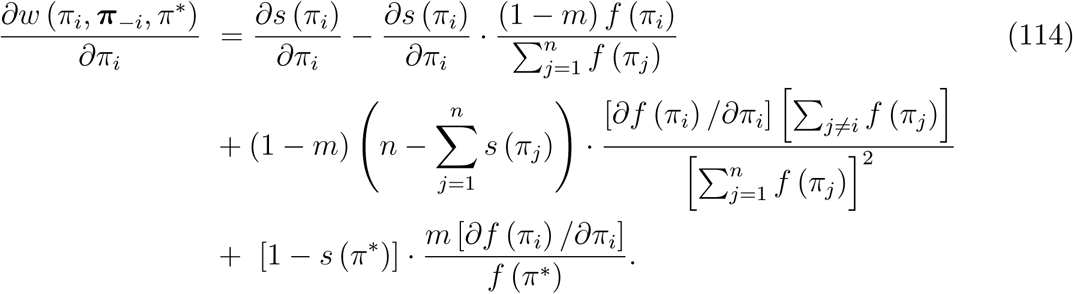

Upon simplification, we obtain:

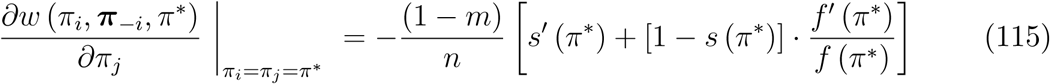

and

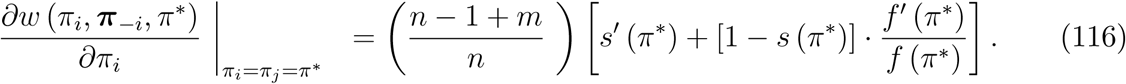

Hence:

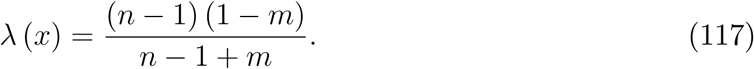

For *r* (*x, x*) the recursion equation writes

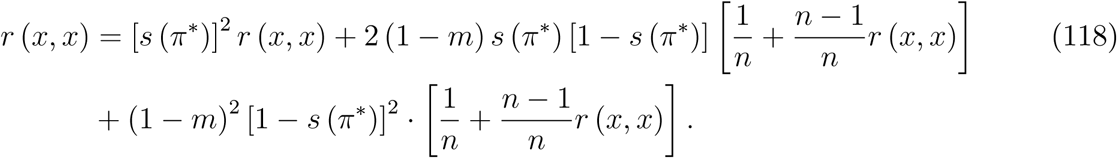

The first term on the right-hand side corresponds to the event that both the individual at hand and the randomly drawn neighbor have been loyal to their parents, where the neighbor’s parent belongs to the individual’s lineage with probability *r* (*x, x*). The second term on the right hand side corresponds to the event that either the individual at hand was loyal to its parent but the randomly drawn neighbor was not loyal to its parent, or the other way around. In this case, there is a probability 1*/n* that the non-loyal child acquired its trait from the loyal child’s parent, in which case they both belong to the same local lineage with certainty, while with the complementary probability this did not happen, in which case the probability that the randomly neighbor belongs to the same local lineage is *r* (*x, x*). The third term on the right hand side corresponds to the event that neither the individual at hand nor the randomly drawn neighbor were loyal to their parents but both of them acquired their trait from someone in the island. In this case, there is a probability 1*/n* that they acquired their type from the same adult, in which case they belong to the same local lineage with certainty; with the complementary probability they have different cultural parents, in which case the probability that the randomly drawn neighbor belongs to the same local lineage as the individual at hand is *r* (*x, x*). We note that the equation simplifies to

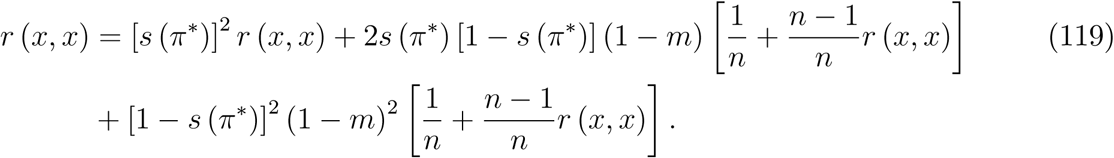

The expression in the text obtains upon observing that this equation is identical to the one in (97).

## Footnotes

1 Throughout this paper we use concepts and terminology that are standard in economics, and model behavior as a choice of action (or stream of actions) from a set of feasible actions, where this choice is guided by a striving to maximize some goal (utility) function. The utility function together with the information and the constraints imposed by the environment are thus what biologists would call the proximate causes driving behavior. Furthermore, by contrast to the evolutionary biology literature where the terms “altruism” and “spite” are used to refer to the fitness consequences of a behavior on the actor and others, in economics they are used to describe the proximate causes behind behaviors. Thus, in economics, an individual who has a utility function which puts a positive weight on another individual’s material well-being is altruistic; and an individual who has a utility function which puts a negative weight on another individual’s material well-being is spiteful. For further discussion of the meaning of these terms in different academic disciplines, we refer to West, Griffin, and Gardner (2007) and Bshary and Bergmüller (2008).

2 A necessary and sufficient condition for kin selection to take place is that an evolving genetic (or cultural) trait tends to more strongly affect the survival and/or reproduction of individuals who are genetically (or culturally) related to the actor than under uniform random matching (Michod, 1982, p.20). This is true whether or not relatives recognize each other. Our analysis will be based on the assumption that they cannot recognize each other.

3 In the economics literature it has been shown that the following conditions are sufficient (and necessary except in knife-edge settings) for populations of self-interested individuals to resist invasion by non-self-interested individuals: *(i)* the population is very large and homogeneous (no subdivision by sex, age, size, etc.) and reproduction is clonal, *(ii)* interacting individuals do not know each other’s preferences but have statistically correct beliefs, and *(iii)* interactions are uniformly random in the population, in the sense that each encounter is just as likely (see Ok and Vega-Redondo, 2001, Dekel, Ely, and Yilankaya, 2007).

4 By contrast to the present model, assortativity was there modeled as an abstract function that maps the distribution of traits in the population to probabilities governing the matching of interacting individuals. This formalization of assortativity was pioneered in economics by Bergstrom (1995, 2003), who focused on strategy evolution; see also Bowles and Gintis (1998), as well as Alger and Weibull (2010, 2012) for analyses of preference evolution under complete information. This formalization of assortativity, which implicitly assumes marginal effects of traits on fitness, goes back to Hamilton (1971); Michod and Hamilton (1980) discuss how different formalizations of assortativity are equivalent to each other. It should further be noted that Rogers (1994) studied the evolution of time preference in an age-structured population; a setting that allows for kin selection but not kin competition. Finally, alternative models of endogenous assortativity have been proposed by Nax and Rigos (2016), Newton (2017), and Wu (2017, 2019).

5 Technically, we study the limit of uniform random dispersal among finitely many islands as the number of islands tend to infinity.

6 For simplicity, we will henceforth use “material payoff” to refer to “expected material payoff”.

7 More precisely, for an)y *x*_*i*_ ∈ *X* and ***x***_−*i*_ ∈ *X*^*n*−1^, and any bijection *h* : {2, 3, …, *n*} → {2, 3, …, *n*}: *π* (*x*_*i*_, *x*_*h*(2)_, *x*_*h*(3)_, …, *x*_*h*(*n*)_ = *π* (*x*_*i*_, ***x***_−*i*_).

8 Continuity is inessential for much of the analysis. However, it is important for some existence results, and for our stability analysis since we there invoke Berge’s maximum theorem. (A form of upper semi-continuity would be sufficient for these results, but such a generalization does not seem to be of primary interest here.)

9 Individual fitness is thus assumed to be expressible in terms of expected material payoff only, and we therefore neglect effects of variance in payoff (see Appendix 6.1 and 6.2 for a justification).

10 By continuity of the utility function and compactness of the set *X*, the set of maximands in (1) is non-empty and compact (by Weierstrass’ maximum theorem). Moreover, by Berge’s maximum theorem, the set of maximands also define an upper hemi-continuous correspondence. By Kakutani’s fixed-point theorem, the set *X*_*u*_ is therefore non-empty (and compact) if the function *u* is also quasi-concave in its first argument (the player’s own strategy).

11 Extinction is defined as the event that no individual in the population is of the mutant type.

12 Even if locally extinct, members of the individual’s lineage may still live on other islands. Moreover, some lineage members may later move to the mutant’s native island. However, the probability that this event occurs in finite time is zero.

13 Note that because the analysis focuses on the fitness of rare mutants in an otherwise homogenous population, our assumption that the fitness function depends on the average material payoff in the population at large, and not on the distribution of the material payoffs therein, is innocuous.

14 This would be true even if migration probabilities were endogenous, as long as migration entails some cost (for literature with endogenous dispersal decisions, see, e.g., Clobert et al., 2001, Frank, 1998, and Rousset, 2004, and Hartl and Clark, 2007). The model by Newton (2017) can be interpreted as having costless migration.

15 In Lehmann, Alger, and Weibull (2015) we proved this result for scenarios where new islands can be colonized only by singleton mutants. Lehmann et al. (2016, eqs. (14)-(16)) extended that result to allow for scenarios in which multiple offspring from the same group can reproduce in the same non-natal island.

16 The reader may worry about model robustness at this point. For if the total population is large but finite, then the probability is not zero, but small and positive, that there will at least one mutant in a given resident’s island. However, for sufficiently large populations (with fixed island size *n*), the probability that a mutant is present in a resident’s island is so small that, by upper hemi-continuity of the best-reply correspondence of any *u* ∈ ℱ, the set of Nash equilibrium strategies for the residents when mutants are very rare in their islands, can be kept within arbitrarily small distance from the set *X*_*u*_.

17 Note that the proposition implies that strategy-committed types that are uninvadable by other strategy-committed types, are uninvadable by all preference types. Indeed, if 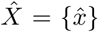, then *u* ∈ ℱ is uninvadable if and only if 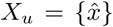. Moreover, this is true even if the residents would have preferences that do not entail commitment to a particular strategy, as long as these preferences induce them to play the uninvadable strategy in residential Nash equilibrium.

18 Obtaining exact expressions for the matching probabilities is typically hard. However, their values can be approximated (see Appendix 6.7 for an approximation method).

19 These preferences are reminiscent of *homo moralis* preferences (Alger and Weibull, 2013, 2016). However, there are two important distinctions. First, here the utility function is defined for a certain reference strategy. Second, the weights attached to the different terms in the utility function depend on the strategy used by the individual at hand.

20 The uni-dimensionality assumption is inessential. All analysis can be carried out in terms of gradients, but with little gain in terms of qualitative insight. For brevity and clarity, we therefore stay with the unidimensional case.

21 First-order conditions like equation (14) apply more generally to traits if lineage fitness and individual fitness are differentiable in trait values. The aforementioned evolutionary dynamics literature focuses on the evolution of phenotypes—the composite of an organism’s characteristics–thus subsuming virtually any heritable trait and can be applied to essentially any demographic scenario (see Rousset, 2004, for general results).

22 This formalization of weak selection corresponds to what Wild and Traulsen (2007) call *w*-weak selection.

23 To be more explicit about this statement, we note that first-order conditions similar to the one in (23) appear elsewhere in the evolutionary biology literature, but then under the form 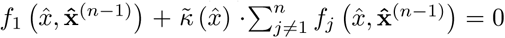, where *f* is the fecundity of an individual (see Lehmann and Rousset, 2010, Akçay and van Cleve, 2012, Van Cleve, 2015, Dos-Santos and Peña, 2017). We would obtain the exact same expression if in our model fitness depended solely on fecundity, since then derivatives of fecundity with respect to material payoffs would cancel from first-order conditions (to see this, set survival to zero in the fitness function in (26) in Section 4). Our model generalizes previous models, since Proposition 5 applies regardless of whether fitness depends only on fecundity, or also on individual and/or group survival (see Section 4 for examples of fitness functions), and it makes explicit the role of the coefficient of fitness interdependence. Further, it demonstrates that even if 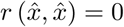, the substitution rate 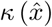 can be substantial depending on the scenario (see, in particular, the examples in Section 4.2). As such, our results unify and extend previous ones of the evolutionary biology literature.

24 The early evolutionary biology literature, which considered traits affecting environmentally induced group extinction (e.g., Eshel, 1972, Aoki, 1982), produced the opposite insight, namely, that pro-sociality at the fecundity level (the equivalent of 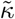 referred to in Footnote 23) is a decreasing function of *n* (see also Lehmann and Rousset, 2010.

25 The analytical models of Bowles (2006, 2009) for the evolution of “parochial altruism” are also close to our scenario with wars; in particular, the expected number of groups 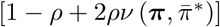 to which a focal group has access for reproduction after warfare also appears in Bowles’s formalization. However, since in his model there are no explicit assumptions that allow to close the lifecycle, it is impossible to derive the explicit values of *λ*_0_, *r*_0_, and *κ*_0_ for his model.

26 A host of other hypotheses about group strength could be explored, see, e.g., Konrad (2014) and the references therein.

27 In the economics literature on cultural transmission of traits, a commonly used model is that of Bisin and Verdier (2001). Like in our model, in Bisin and Verdier (2001) each grown-up has exactly one child, and each child inherits its parent’s trait with some probability, and otherwise it inherits the trait of another grown-up in the population. By contrast to our model, the population is not structured into islands, and there is no strategic interaction between individuals. Furthermore, in their model a parent cares about whether her child has the same trait as her, but not about whether the child inherited this trait from the parent or from someone else. Denoting by *q*_*θ*_ the population share of individuals with trait *θ* in the population, and by *s* (*q*_*θ*_) the probability that a child inherits its trait vertically from its parent, their equation (1) says that the unique child of a parent with trait *θ* acquires trait *θ* with probability *s* (*q*_*θ*_)+ [1 − *s* (*q*_*θ*_)] · *q*_*θ*_. Thus, in the their model it is only the frequency of the trait that determines the transmission probability; in our model the attractiveness of a trait also plays a role.

28 In evolutionary biology, the same expression as the right hand side of (42) was obtained for the case of no cultural loyalty as a marginal fecundity cost to benefit threshold ratio under which the mutant is favored in a public good game (eq. 26 of Lehmann, Foster, and Feldman, 2008).

29 To be more specific, and using Scenario A to illustrate this point, if the mappings *f* and *s* in (26) change, while the migration probability *m* as well as the function *w* remain unchanged, then the weights attached to the components in *v*^0^ remain unchanged.

